# A benchmark of muscle models to length changes great and small

**DOI:** 10.1101/2024.07.26.605117

**Authors:** Matthew Millard, Norman Stutzig, Jörg Fehr, Tobias Siebert

## Abstract

Digital human body models are used to simulate injuries that occur as a result of vehicle collisions, vibration, sports, and falls. Given enough time the body’s musculature can generate force, affect the body’s movements, and change the risk of some injuries. The finite-element code LS-DYNA is often used to simulate the movements and injuries sustained by the digital human body models as a result of an accident. In this work, we evaluate the accuracy of the three muscle models in LS-DYNA (MAT_156, EHTMM, and the VEXAT) when simulating a range of experiments performed on isolated muscle: force-length-velocity experiments on maximally and sub-maximally stimulated muscle, active-lengthening experiments, and vibration experiments. The force-length-velocity experiments are included because these conditions are typical of the muscle activity that precedes an accident, while the active-lengthening and vibration experiments mimic conditions that can cause injury. The three models perform similarly during the maximally and sub-maximally activated force-length-velocity experiments, but noticeably differ in response to the active-lengthening and vibration experiments. The VEXAT model is able to generate the enhanced forces of biological muscle during active lengthening, while both the MAT_156 and EHTMM produce too little force. In response to vibration, the stiffness and damping of the VEXAT model closely follows the experimental data while the MAT_156 and EHTMM models differ substantially. The accuracy of the VEXAT model comes from two additional mechanical structures that are missing in the MAT_156 and EHTMM models: viscoelastic cross-bridges, and an active titin filament. To help others build on our work we have made our benchmark simulations and model code publicly available.

## 1 Introduction

Digital human body models (HBM) are used to evaluate the risk of injury during low-velocity vehicle collisions [1], [2], from exposure to vibration [3]–[5], and as a result of athletic accidents [6], [7]. Simulating injury-causing scenarios is challenging because the musculature of the body may have time to activate [8], altering the ensuing movement [9], [10], and affect the risk of some types of injury. When activated, muscle develops tension and its mechanical properties change: active muscle can generate large forces in response to modest stretches [11]–[13], and the stiffness and damping (impedance) of active muscle can increase substantially [14]. Unfortunately, simulations that involve active-lengthening or the vibration of muscle should be approached with caution: few muscle models have been evaluated for accuracy during either active lengthening or vibration.

Nearly all digital HBMs with active musculature use the Hill-type muscle models [1], [15]–[19] despite the limitations of this formulation. Ritchie and Wilkie [20] derived the Hill-type muscle model in 1958 with the aim of simulating four experimentally observed phenomena: the variation of isometric force with the length of the contractile-element (CE), the variation of CE force with velocity, the time-dynamics of muscle force during activation and deactivation, and the interaction between the CE and a serially-connected elastic tendon. Within these four experimental phenomena Hill-type muscle models have limitations. Most Hill-type muscle models are able to capture the force-length-velocity properties of maximally activated muscle but not of sub-maximally activated muscle [21], [22]. Few Hill-type muscle models [23], [24] have been evaluated in the context of active-lengthening, particularly at long CE lengths [11], though this comprises half of the force-velocity relation.

Models used to simulate injury are typically evaluated by simulating an entire musculoskeletal model rather than evaluating the individual components of the model. While it is necessary to examine the accuracy of a musculoskeletal model to simulate a particular injury [5], [8], [19], [25], these simulations offer little insight into whether individual muscles are being simulated accurately because the corresponding experimental data is necessarily incomplete: it is not possible to measure the three-dimensional boundary conditions and forces of the body’s musculature in a living person. Experiments on isolated muscle, in contrast, make it possible to control the boundary conditions and measure the forces developed by muscle.

While the literature has many simulations of classic muscle physiology experiments — activation dynamics [26], force-length-velocity relations [27], [28], force-depression and enhancement [29] — there are comparatively few works that include experiments that are relevant for active-lengthening injury [30], [31] and the vibration response [14] of muscle. There are also relatively few works that examine the muscle models [32] available in LS-DYNA, a finite-element (FE) code that is commonly used to simulate digital HBMs. Our recent simulation study [33] shows that there are reasons to be concerned about the accuracy of muscle models during simulations of injury and vibration: the simulated forces developed during modest [11] and extreme lengthening [12] are lower than experimental data, and the response of the model to vibration is more damped than biological muscle [14]. There are a wide variety of Hill-type muscle model formulations, and so, it is not clear how well the muscle models implemented in LS-DYNA will fare when simulating experiments that examine active-lengthening [11], and frequency-response^1^ [14] of muscle.

In this work, we extend the work of Kleinbach et al. [32] by assessing the accuracy of three muscle models in LS-DYNA [34] by simulating four different types of experiment: isometric force-length experiments, force-velocity experiments at short CE lengths, active-lengthening experiments at long CE lengths, and the response of the muscle to vibration. The models range in structural complexity, from the basic Hill model provided by LS-DYNA [35] (MAT_156), to the extended Hill-type muscle model (EHTMM) that includes a viscoelastic tendon [27], [32], [36], and, finally, to a recently introduced model [33], [37] that includes a viscoelastic crossbridge and active titin elements (VEXAT). We simulate experiments to illustrate both the strengths and weaknesses of muscle models when simulating maximal and submaximal force-length, and force-velocity experiments. In addition, we include experiments that are specifically relevant for the simulation of injury: active lengthening on the descending limb and the response of muscle to vibration. Our analysis focuses specifically on the muscle models that are available in LS-DYNA [34] because LS-DYNA is commonly used for crash simulation and for the simulation digital HBM in general. So that others can extend our work we have made the LS-DYNA implementation of the VEXAT model^2^ and benchmarking simulations^3^ available online.

## 2 Models

Our benchmark simulations evaluate the responses of three different lumped-parameter muscle models in LS-DYNA [34]: MAT_156 [35], EHTMM [27], [32], [36], and the VEXAT [33] muscle model. These models use a simplified geometric representation (Fig. 1A) of the muscle-tendon complexes where all fibers in the CE are lumped to one side and are assumed to be identical and act in series with an elastic tendon. The geometric model used for pennated muscle has an overall path length (*𝓁* ^P^) given by

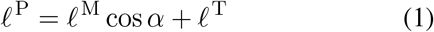

where *𝓁* ^M^ is the length of the CE, *α* is the angle between the CE and the tendon (Fig. 1A, bottom), and *𝓁* ^T^ is the length of the tendon. To mimic the constant volume property of muscle [38], the muscle is assumed to have a fixed depth and the pennation angle *α* is varied such that height of the CE

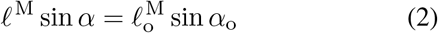

remains constant, where 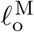 is the length of the CE at which the largest force is developed (Fig. 1C), and *α*_o_ is the pennation of the CE at 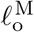. Where the VEXAT model includes a pennation model [33] (Fig. 1A, bottom), both LS-DYNA’s MAT_156 and the EHTMM can only represent non-pennated muscles (Fig. 1A, top). This difference in geometric modeling is of little consequence for the benchmark simulations that follow because the muscles simulated have small values of *α*_o_^4^.

**Figure 1:**
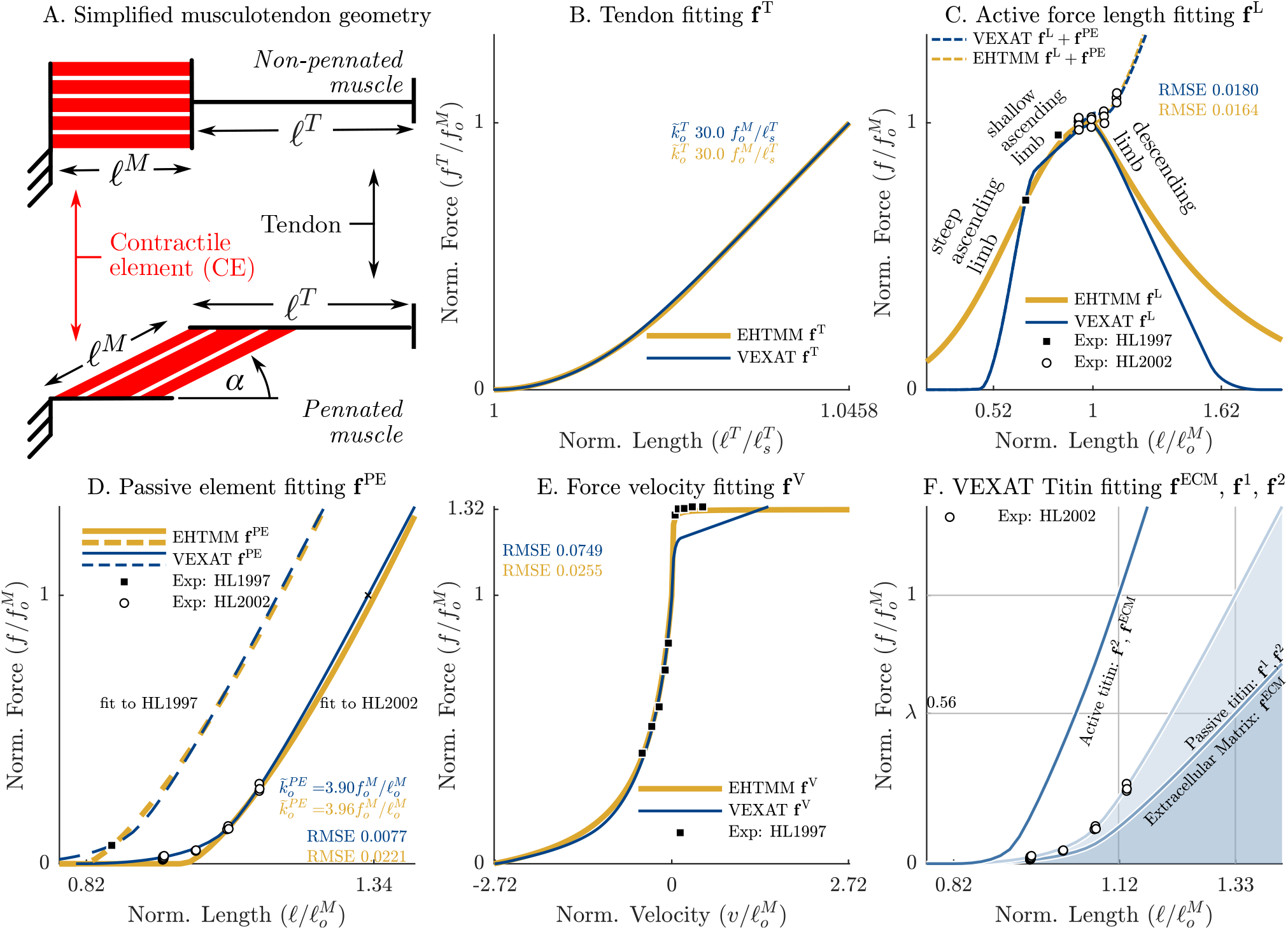
The models evaluated in this work represent muscle geometrically as a one-dimensional cable that has a contractile-element (CE) in series with a tendon (A). The CE may act in the same direction as the tendon (A, top), or at an angle (A, bottom) called the pennation angle. To mimic the constant volume property of muscle [38], the angle of a pennated CE is varied to have a constant height which endows the resulting fixed-depth parallelepiped with a constant volume. Muscle and tendon have a number of non-linear characteristics represented by parametric equations in the VEXAT [33] and EHTMM [27], [32], [36] models: the force-length relation of the tendon (B, which has a stiffness of 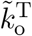 at a tension of 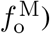, the active-force-length relation of the CE (C), the passive force-length relation of the CE (D, which has a stiffness of 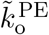 at a passive tension of 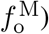, the force-velocity relation of the CE (E). We have set the tabular data used by the MAT_156 to follow the curves of the VEXAT model. The VEXAT model has additional non-linear curves (F) to represent the force-length relations of extracellular matrix (ECM), the proximal segment of titin, and the distal segment of titin. When activated, the proximal segment is approximately fixed and, as a result, the active titin segment appears stiffer when stretched (F). While there are differences between the parametric equations of the EHTMM and the VEXAT models the root-mean-squared-error (RMSE) of these to models relative to the experimental data is similar (B, C, D, and E).

Each of the muscle models is dimensionless but can be scaled to any muscle-tendon complex using its architectural properties: the maximum active isometric force 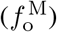, the optimal fiber length 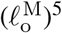, the maximum shortening velocity of the CE 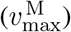, and the slack length of the tendon 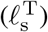. These architectural properties are used to scale thecurves that have been fit to capture experimentally observed relationships: the force-length relation of the tendon [40], [41] (**f** ^T^, Fig. 1B), the active force-length relation [42] (**f** ^L^, Fig. 1C), the passive force-length relation [43] (**f** ^PE^, Fig. 1D), and the force-velocity relation [44] (**f** ^V^, Fig. 1E) of the CE. The VEXAT model [33] further decomposes **f** ^PE^ into the elastic contributions from three smaller structures (Fig. 1F): the extracellular matrix (ECM, 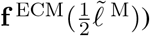, titin’s proximal segment 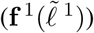, and titin’s distal segment 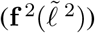. To facilitate scaling, each of these relations are described in terms of normalized length, normalized velocity, and normalized force. Throughout this manuscript we use a tilde to indicate a normalized quantity: within the CE length is normalized by 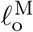, velocity by 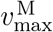, and force by 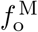 while tendon length is normalized by 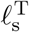 and force by 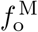. In addition, curves are indicated using bold font, for example, the force-length relation 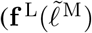 pictured in Fig. 1C). Although the MAT_156, EHTMM, and VEXAT models use different parametric equations for the force-length-velocity curves (Fig. 1B-C), all of these curves use the same normalization factors and have broadly similar shapes. Despite these similarities, each model represents different mechanical structures of a muscle-tendon complex.

LS-DYNA’s MAT_156 includes a stateless two-component model of the CE (Fig. 2A ^6^) and does not include a tendon model. The force (Fig. 2B) developed by MAT_156’s CE is the sum of the passive and active components

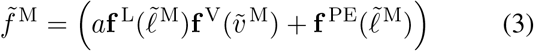

where *a* is a 0-1 quantity that represents the level of chemical activation. The curves used to describe 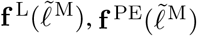, and 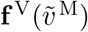 are represented using tabular data that set to the VEXAT model’s curves in this work.

**Figure 2:**
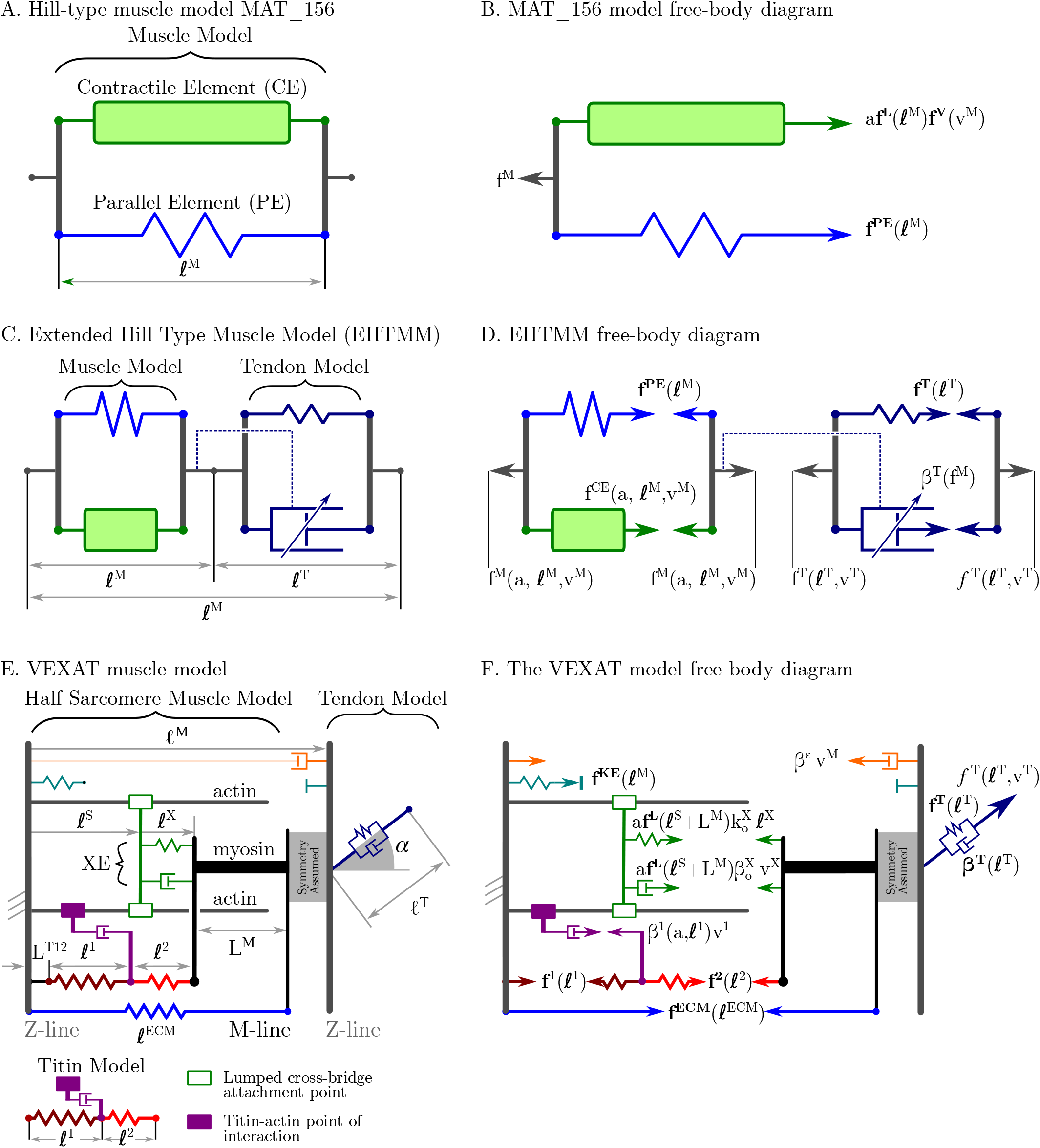
LS-DYNA’s MAT_156 consists of a CE that is in parallel with an elastic element (A), such that the total force developed by the model is the sum of the active and passive elements (B). The EHTMM, formulated by Gü nther et al. [27], extended by Haeufle et al. [36] and implemented in LS-DYNA by Kleinbach et al. [32], is composed of a CE in series with a viscoelastic tendon (C). The CE and tendon are assumed to be in a force equilibrium (D) which Günther et al. [27] solves efficiently by assuming that the tendon damping follows a specific function. The VEXAT model [33] has a three component CE (viscoelastic XE, an active titin model, and a passive ECM) in series with a viscoelastic tendon (E). The XE is the only element capable of doing net positive work (F), the ECM is passive, and the stiffness of the titin element is modified by activation. We model as two serially connected springs that meet at a location that can become bound to actin when the CE is active.

The EHTMM includes a viscoelastic tendon (Fig. 2C), a state *𝓁* ^M^, and a differential equation for *v* ^M^ that can be numerically integrated forward in time to yield the trajectory *𝓁* ^M^(*t*) [27]. The CE of the EHTMM embeds the force-length relation into to Hill’s [44] force-velocity relation

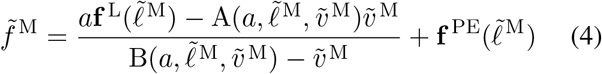

by cleverly formulating the Hill parameters 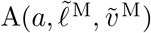 and 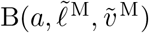 to create a force-length-velocity curve in which 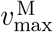 varies with *a* similar to biological muscle [27].However, Eqn. 4 cannot be evaluated directly because *v* ^M^ is unknown. To solve for *v* ^M^, it is assumed that the CE and the tendon are in a force equilibrium

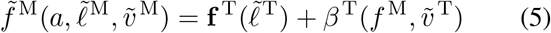

with the viscoelastic tendon (Fig. 2D). To solve for 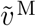, most Hill-type formulations either solve for 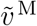 directly and introduce a singularity in the solution [45], [46], or regularize the equation using an additional damping element which results in a nonlinear equation that can only be solved numerically [47]. Instead, Günther et al. [27] assumed that the tendon’s damping force 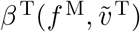 takes a specific form that turns Eqn. 5 into a function that is quadratic in 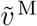, making it possible to efficiently solve Eqn. 5 directly for 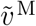. The EHTMM uses power functions to describe **f** ^PE^ and **f** ^T^, exponential functions for **f** ^L^, and hyperbolas for **f** ^V^.

The VEXAT model (Fig. 2E) introduces a lumped viscoelastic crossbridge (XE) as well as two-segment active model of titin [33]. This model has a total of four states: the XE’s attachment position (*𝓁* ^S^) and velocity (*v* ^S^), the length of the proximal segment of titin (*𝓁* ^1^), and the length of the CE (*𝓁* ^M^). The tension

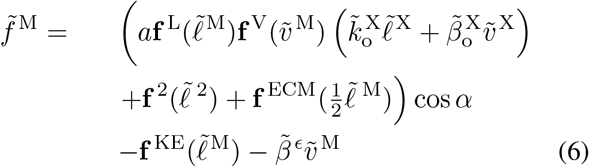

developed by the VEXAT’s CE (Fig. 2F) is dominated by contributions from the XE’s stiffness 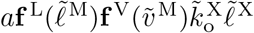 and damping 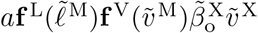, the distal elastic segment of titin 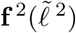, and the elasticity of the ECM 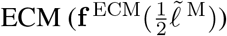. The remaining two terms are in place for practical reasons: 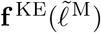 prevents the CE from reaching unrealistically short lengths while 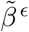 is small damping added for numerical stability. The state derivative of the VEXAT model [33] is evaluated directly by assuming that the proximal 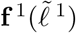 and distal 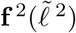 segments are in a force equilibrium, that the CE and tendon are in a force-equilibrium, and such that the XE slowly tracks force of a Hill model (Fig. 2F). The numerous curves in the VEXAT model are implemented as Bézier spines.

## 3 Benchmark Simulations

We have selected four experiments to simulate in order to compare and contrast the MAT_156 [35], EHTMM [27], [32], [36], and VEXAT [33] muscle models: the force-length relation of passive and active muscle [48]–[50], the force-velocity relation on the *ascending limb* [51] of the force-length relation 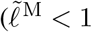 in Fig. 1C), active lengthening on the *descending limb* [11] of the force-length relation 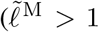 in Fig. 1C), and the impedance-force^7^ relation [14]. The benchmark simulations of force-length and force-velocity relations are included both to serve as a context for later simulations and also so that we can evaluate how the models perform during submaximal contraction. The active-lengthening and impedance benchmark simulations have been included because both of these relations are relevant for the simulation of injury. We have intentionally chosen to simulate experiments using in-situ cat soleus for two reasons: an in-situ preparation is as close to in-vivo conditions as is possible in a lab setting, and there are numerous studies on cat soleus that can be used to both fit and evaluate the models.

### 3.1 Model Fitting

Prior to evaluating the accuracy of the models when simulating the force-length, force-velocity, eccentric, and impedance of muscle we must fit the parameters of each of the models. To simulate these four experiments, we need a total of four cat soleus model variants: a model (HL97) fitted to Herzog and Leonard 1997 [51], a model (HL02) fitted to Herzog and Leonard 2002 [11], and models (K3, K12) to simulate the data from Figures 3 and 12 from Kirsch et al. [14]. To ensure that our simulations are as fair as possible, we have fit the models with two aims in mind: to match the experimental data as closely as possible such that each model has curves 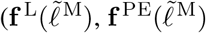, and 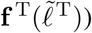 that are as similar as possible. While many parameters are identical between the four cat soleus variants, the differences that exist are primarily in the architectural properties (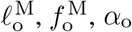, and 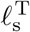) which is expected because these experiments were performed on different specimens.

**Figure 3:**
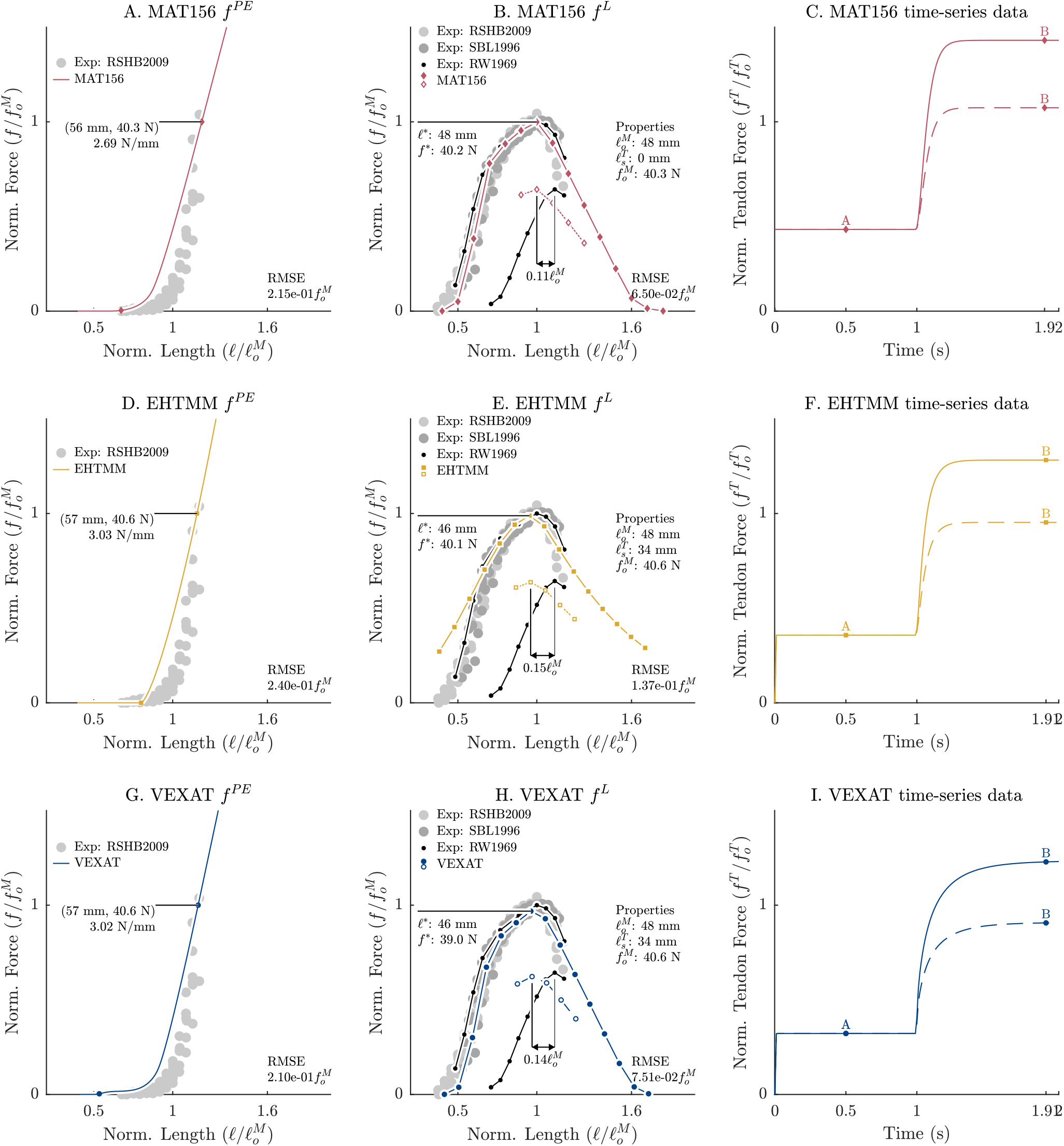
When each of the models is fit using the parameters of HL97, each model has a similar level of accuracy when compared to the testing data: Rode et al. ([50], RSHB2002), Scott et al. ([48], SBL1996), Rack and Westbury ([21], RW1969). The passive force-length relations (A, D, and G) reach the strains and stiffness needed to give all three muscle-tendon complexes the same length and stiffness when 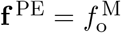. Note that the MAT_156’s CE is more compliant than the other two models because its compliance must match the other two models which have elastic tendons. Each of the models have a maximally active force-length relation (solid lines in B, E, H) that follows the testing data closely, though the EHTMM deviates where 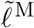 is outside of the range 0.67 − 1.14. The submaximal active force-length relation of each model (dashed lines in B, E, H) has a peak that deviates from the experimental data. The active force-length relations (B, E, and H) are created by performing simulations in which the path length of the muscle is held constant while it is activated (C, F and I). Using a tendon force-length model and a model of the passive force-length relation (created using the ‘A’ measurements in (C)) the CE length and active force is evaluated for the ‘B’ measurements and this is used to form one point in the active force-length relation (B, E, and H). Finally, although all three models use a Zajac [45] activation model the VEXAT model (I) develops force more slowly because of the time-dynamics of its XE.

The parameters for the four different model variants are fitted in four stages: first, active and passive force-length parameters are determined for both HL97 [51] and HL02 [11]; second, force-velocity parameters for all models are set using Herzog and Leonard 1997 [51]; third, active titin model parameters are set for all VEXAT model variants using Herzog and Leonard 2002 [11]; finally, the stiffness and damping of XE is set for the VEXAT model variants K3 and K12 using the data in Figures 3 and 12 of Kirsch et al. [14], respectively. Since each of these experiments measures only a few properties each model variant uses parameters fitted to other studies: the passive force-length, active force-length, and titin properties from HL02 are also used in model variants K3 and K12; the force-velocity properties of HL97 are used for all other model variants; the XE parameters for K3 is applied to the VEXAT model variants HL97, and HL02. Although it is unsatisfying to require data from many different experiments this is necessary: no single experiment in the literature contains all of the information required to fit all of the parameters of a muscle model. We do not expect the heterogeneous mix of parameters to introduce much error since many characteristics are similar from one muscle to the next when CE lengths are normalized by 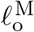 [54], CE velocities by 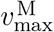 [55], forces by 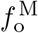 [56], and tendon lengths by 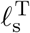. In the following paragraphs we present an overview of the fitting process we have used for this work while the technical details can be found in Appendix A.

The force-length properties of all of the model variants are set using the data from Herzog and Leonard 1997 [51] and 2002 [11]. Both of these studies [11], [51] include ramp-lengthening and shortening trials which inherently include a sampling of the passive force-length relation, the active force-length relation and the force-velocity relation. However, there are 3 experimental parameters that are either missing or are uncertain in each study: the optimal CE length 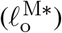, the maximum isometric force 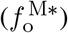, the maximum shortening velocity 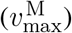, and the path length of the muscle that corresponds to the reference length (*𝓁* ^R*^) of 0mm. Since the VEXAT model’s active and passive force-length relations require relatively few parameters, we first solve for the experimental parameters 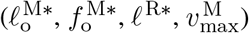, passive force-length parameters (Δ^*^ and *s*^*^ which shift and scale the 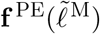 of the VEXAT model) that best fit of Herzog and Leonard 1997 [51] and 2002 [11] simultaneously (see Appendix A for details).

Next, the shape of passive and active force-length relations of the EHTMM are fitted to the data [11], [51]. During the fitting process the values for 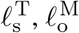 and 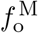 of EHTMM model are set to 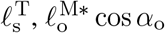 and 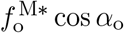 from the VEXAT model so that both of these models are as similar as possible when evaluated in the direction of the tendon. In addition, the EHTMM uses the starting path length (*𝓁* ^R*^) identified using the VEXAT model so that both models are simulated using the same boundary conditions (see Appendix A for details). The fitting process produces a set of passive and active force-length parameters for the VEXAT and EHTMM models for HL97 (see Table 1) and HL02 (see Table 2) variants.

**Table 1:**
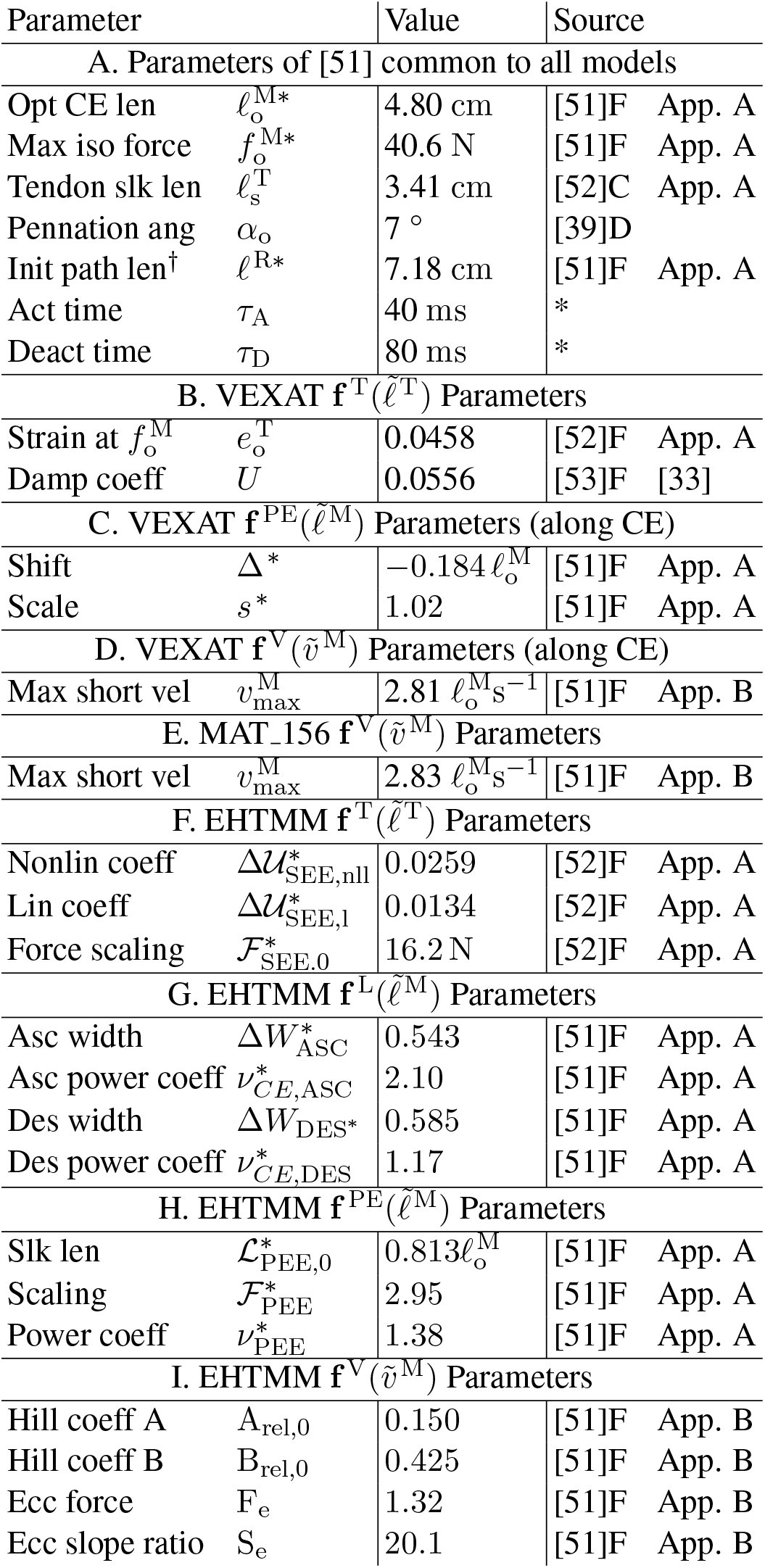
The force-length-velocity model parameters applied to model variant HL97. The following short forms are used in the interest of space: optimal (opt), length (len), maximum (max), isometric (iso), slack (slk), angle (ang), velocity (vel), initial (init), activation (act), deactivation (deact), coefficient (coeff), ascending limb of the force-length relation (asc), descending limb of the force-length relation (des), nonlinear (nonlin), linear (lin), and eccentric (ecc). The source column begins with a reference for the parameter and is followed by a letter to indicate how the data was used: ‘D’ for directly used, ‘F’ for fit, and ‘C’ for calculated. Parameters *τ*_A_ and *τ*_D_ (*) have not been fitted because simulations are evaluated under constant activation.

**Table 2:** The force-length-velocity parameters used for model variants HL02, K3, and K12. All of the conventions from Table 1 are used in this table. In addition, parameters that differ from Table 1 are highlighted in gray. Since these parameters are for a different cat soleus than Table 1 the architectural properties differ, as do the properties of the tendon and the passive elasticity of the CE.

| Parameter |  | Value | Source |
| --- | --- | --- | --- |
| A. Parameters of [11] common to all models |  |  |  |
| Opt CE len | $\ell_o^{M*}$ | 4.90 cm | [51]F App. A |
| Max iso force | $f_o^{M*}$ | 21.6 N | [51]F App. A |
| Tendon slk len | $\ell_s^T$ | 3.45 cm | [52]C App. A |
| Pennation ang | $\alpha_o$ | 7 ° | [39]D |
| Init path len<sup>†</sup> | $\ell^{R*}$ | 8.17 cm | [51]F App. A |
| Act time | $\tau_A$ | 40 ms | * |
| Deact time | $\tau_D$ | 80 ms | * |
| B. VEXAT $\mathbf{f}^T(\tilde{\ell}^T)$ Parameters | | | |
| Strain at $f_o^M$ | $e_o^T$ | 0.0458 | [52]F App. A |
| Damp coeff | $U$ | 0.0556 | [53]F [33] |
| C. VEXAT $\mathbf{f}^{PE}(\tilde{\ell}^M)$ Parameters (along CE) | | | |
| Shift | $\Delta^*$ | $-0.0172 \ell_o^M$ | [51]F App. A |
| Scale | $s^*$ | 1.02 | [51]F App. A |
| D. VEXAT $\mathbf{f}^V(\tilde{v}^M)$ Parameters (along CE) | | | |
| Max short vel | $v_{\max}^M$ | $2.72 \ell_o^M s^{-1}$ | [51]F App. B |
| E. MAT_156 $\mathbf{f}^V(\tilde{v}^M)$ Parameters | | | |
| Max short vel | $v_{\max}^M$ | $2.74 \ell_o^M s^{-1}$ | [51]F App. B |
| F. EHTMM $\mathbf{f}^T(\tilde{\ell}^T)$ Parameters | | | |
| Nonlin coeff | $\Delta \mathcal{U}_{SEE,nll}^*$ | 0.0259 | [52]F App. A |
| Lin coeff | $\Delta \mathcal{U}_{SEE,l}^*$ | 0.0134 | [52]F App. A |
| Force scaling | $\mathcal{F}_{SEE,0}^*$ | 8.62 N | [52]F App. A |
| G. EHTMM $\mathbf{f}^L(\tilde{\ell}^M)$ Parameters | | | |
| Asc width | $\Delta W_{ASC}^*$ | 0.545 | [51]F App. A |
| Asc power coeff | $\nu_{CE,ASC}^*$ | 2.09 | [51]F App. A |
| Des width | $\Delta W_{DES}^*$ | 0.585 | [51]F App. A |
| Des power coeff | $\nu_{CE,DES}^*$ | 1.17 | [51]F App. A |
| H. EHTMM $\mathbf{f}^{PE}(\tilde{\ell}^M)$ Parameters | | | |
| Slk len | $\mathcal{L}_{PEE,0}^*$ | $0.998 \ell_o^M$ | [51]F App. A |
| Scaling | $\mathcal{F}_{PEE}^*$ | 2.07 | [51]F App. A |
| Power coeff | $\nu_{PEE}^*$ | 1.36 | [51]F App. A |
| I. EHTMM $\mathbf{f}^V(\tilde{v}^M)$ Parameters | | | |
| Hill coeff A | $A_{rel,0}$ | 0.153 | [51]F App. B |
| Hill coeff B | $B_{rel,0}$ | 0.418 | [51]F App. B |
| Ecc force | $F_e$ | 1.32 | [51]F App. B |
| Ecc slope ratio | $S_e$ | 20.1 | [51]F App. B |

Now that we have solved for most of the architectural properties (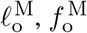, and 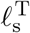) and the force-length relations 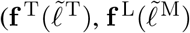, and 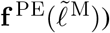 of both HL97 and HL02 we can fit the force-velocity relation to Herzog and Leonard 1997 [51]. As before, we fit the underlying parametric curves of the VEXAT and EHTMM to the experimental data [51], and construct the force-velocity relation of MAT_156 by numerically sampling the projection of the VEXAT model’s force-velocity curve in the direction of the tendon (see Appendix B for details). The fitted **f** ^V^ of all three models has the same maximum shortening velocity in the direction of the tendon (see Table 1D-E, Table 2D-E, and Appendix B) and closely follow the experimental data [51] during shortening (Fig. 1E). The eccentric side of the VEXAT model’s **f** ^V^ produces less force than the experimental data to make room for the force contribution from the active-titin element, which is not included in the **f** ^V^ curve of the VEXAT model.

With the force-length-velocity parameters of all three models fit, we can turn our attention to fitting the active-titin and XE viscoelasticity parameters of the VEXAT model. The VEXAT’s active-titin model includes 12 parameters (see Table 1H of [33]) most of which are related to the geometry of the titin segment and are fixed. There are 2 parameters that we adjust to more accurately simulate the tension developed during active lengthening in Herzog and Leonard’s 2002 [11] study: Q, the point within titin’s PEVK segment that attaches to actin (as Q increases from 0 to 1 the resting length of 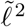 becomes shorter and stiffer, see Fig. 2E), and 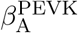, the maximum active damping that is applied between the PEVK segment and actin during active lengthening (as 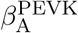 increases, the maximum value that 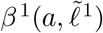 can reach increases, see Fig. 2F). The error used to fit Q is calculated by simulating the 9mm active lengthening trial at 9mm s^−1^ (see Figure 7B of [11]) and subtracting the peak tension developed by the model from the 36.6N peak measured force. Since it is time consuming to evaluate this error, we used the bisection method to solve for the value of Q = 0.593 that resulted in the best agreement with the 9mm trial in Figure 7B [11]. The second active-titin parameter 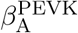 is fit by minimizing the squared error between the force generated by the model and the data ([11], 9mm trial in Figure 7B) at 10 evenly spaced samples during the 5 seconds after the ramp length-change ends. As with the active-titin parameters we used the bisection method to solve for the value 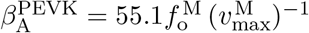.

The values of the maximum active normalized stiffness 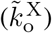 and damping 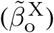 of the XE that best fit Figure 3 and Figure 12 of Kirsch et al. [14] were set to the values that appear in Appendix 2, Table 2 of Millard et al. [33]. The gain and phase profiles from Kirsch et al. [14] and a linearized version of the VEXAT model are used to solve for 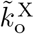 and 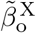 under the assumption that XE remains bound to actin. During simulation, the XE is not perfectly bound to actin even during full activation, and so, this method of fitting 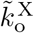 and 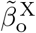 results in a model that will be a bit less stiff than desired. We have set VEXAT model variants HL97, HL02, K12 to the values of 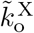 and 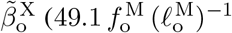 and 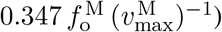 for Figure 12 of Kirsch et al. [14]. Model variant K3 has the higher stiffness 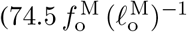 and damping 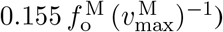 of the specimen illustrated in Figure 3 of Kirsch et al. [14].

### 3.2 Isometric active and passive force-length relations

Although it is frequently assumed that Hill-type muscle models can reproduce the force-length [42], [43] relation, a few details are often overlooked. The shape of the force-length relation [42], [57] of whole muscle may differ [58] from the theoretical model derived from the sliding filament theory [59] because the geometric path of the fibers in whole muscle can differ from that of a scaled sarcomere. In addition, the location of peak isometric force is known to shift to longer lengths during submaximal activation [21], [60]. Since we have fit the shape of the passive and active force-length relations to Herzog and Leonard 1997 [51] and 2002 [11], we evaluate each model against three different data sets [21], [50], [52] one of which also includes submaximal activation [21] trials.

To evaluate the models, we simulate the experiments that are typically used to measure the passive and active force-length relations experimentally. The passive force-length relation is derived by simulating each muscle as it is passively stretched. Next, the model is simulated isometrically beginning from a passive state and ending with a sustained activation at a series of path lengths to sample the force-length relation. Due to activation dynamics and tendon elasticity, the active force of each muscle is sampled after it has been activated long enough to converge to its final value. Finally, the active force is evaluated by subtracting off the passive-force that corresponds to the final CE length: we cannot use the initial passive force since this may differ from the final passive force of the CE due to the elasticity of the tendon [50]. In an experiment this last step can only be done if the length of the CE or tendon is measured as done by Scott et al. [48].

All three of the models are able to follow the fitting data and each other closely (Fig. 1B-D) and provide similar levels of error when compared to the testing data (Fig. 3) for both maximal and submaximal activation. The differences that arise are mostly due to the parametric curves used to define 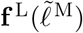 for the EHTMM model: for 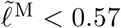 and 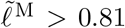 the piece-wise continuous Gaussian function is not able to closely follow the data of Scott et al. [48] and leads to a higher RMSE than the other models. None of the models show a shift in the peak of the active force-length relation with submaximal activation (Fig. 3B,E, and H). This perhaps should not be surprising, as none of the models has a mechanism to shift the active force-length relation with submaximal activation.

### 3.3 Active shortening and lengthening on the ascending limb

While Hill’s force-velocity relation [44] is embedded in the three models evaluated, this alone is not sufficient to guarantee that each model can capture the variation of muscle force with velocity. First, submaximal shortening is often accompanied by a reduction in the maximum shortening velocity that is not captured in the original formulation of Hill’s force-velocity relation [44]. Next, Hill’s force-velocity relation [44] only specifies the change in force during shortening at a specific length at an instant in time. Since experimental methods to measure the force-velocity relation take time, there are time dynamics associated with these experiments that are also not captured by the force-velocity relation. Finally, while Hill’s force-velocity relation [44] has robustly predicted the tension developed during shortening, there is no equivalently consistent model of active lengthening.

To test the models, we simulate Herzog and Leonard’s 1997 [51] experiment in which maximally activated cat soleus undergoes a series of shortening and lengthening trials that all end at a reference length of 0mm which corresponds to *𝓁* ^R*^ in (Table 1A). To simulate the experiment, we digitized both the force and length profiles ([51], Figure 1A) and configured each model to use the HL97 parameters (Table 1). The shortening trials begin with each model in a passive state and with a path length of *𝓁* ^R*^ +4mm. After 1s the model is activated, shortening begins at 1.57s and proceeds at the rate (−2.6, -4.9, -9.8, -16.0, and -23.5 mm s^−1^) calculated from our digitized data ([51], Figure 1A) until the reference length of 0mm is reached. From this point on the length of the model is held fixed until a time of 4.1s to be consistent with the experiment [51]. The lengthening trials are similar except the initial length is *𝓁* ^R*^ − 4mm and the model is lengthened at the rates indicated from our digitized data (2.4, 4.7, 8.8, 13.2, and 21.5 mm s^−1^) until the reference length of 0mm is reached.

The data from these 10 simulations are next transformed into 10 discrete points

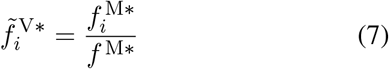

on the force-velocity relation using the force 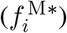 measured during the length change, the isometric force (*f* ^M*^), and the normalized rate 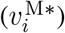 of length change

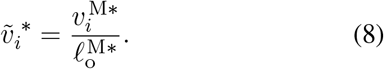

We compare the simulated normalized force-velocity relation to separate testing data that we have manually digitized: Figure 4A^8^ of Scott et al. [48], Figure 8A^9^ from Brown et al. [49], and Figure 5^10^ from Joyce and Rack [61]. We make this comparison using a normalized force-velocity plot to minimize differences between specimens. In addition, we evaluate the root-mean-squared-error (RMSE) between the simulated and measured time-series data in two phases: during the length-change, and during the time period after the length-change has been completed.

**Figure 4:**
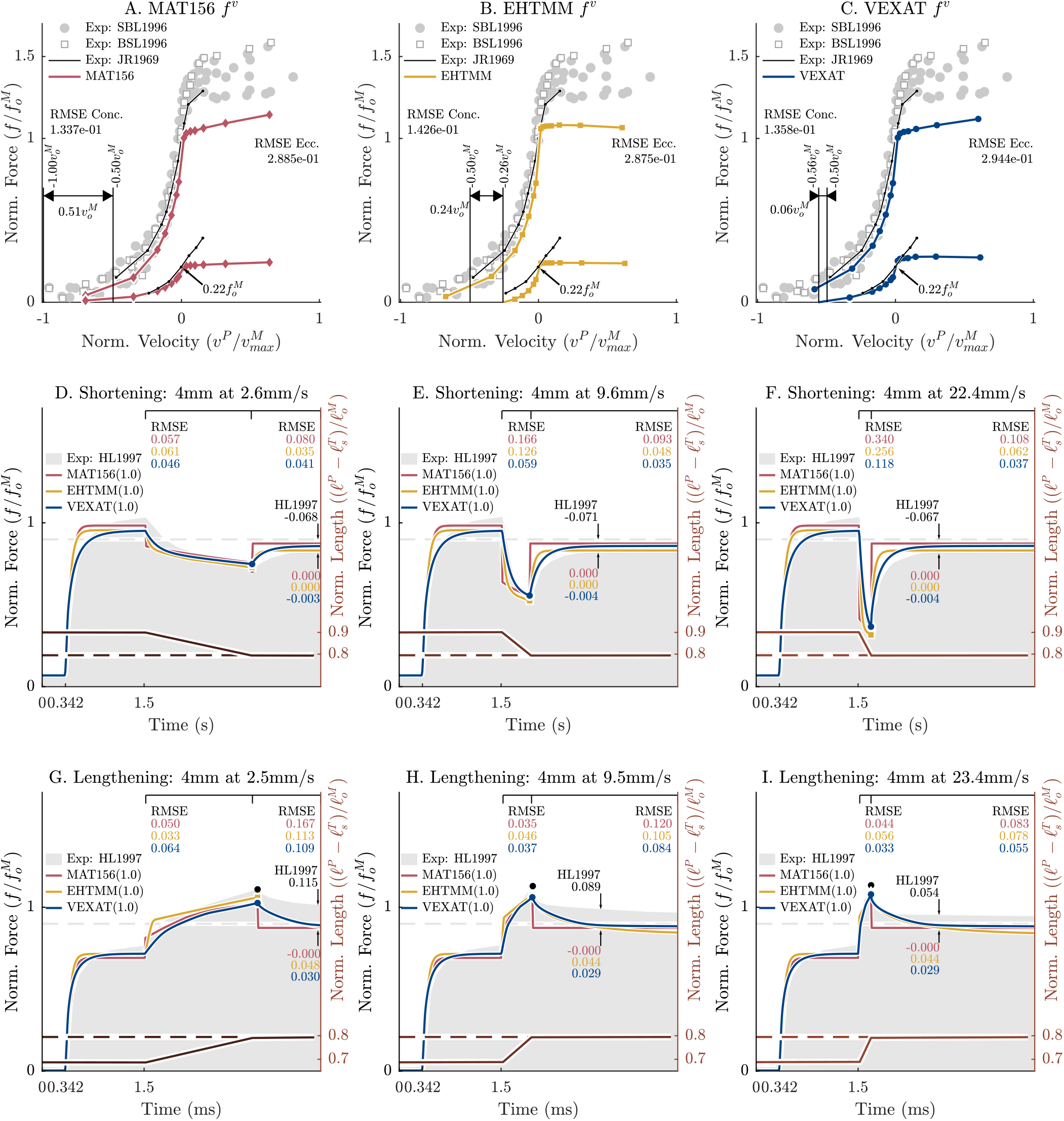
When the force-velocity relation is extracted from isokinetic simulations of each model under maximal activation (A, B, and C) the results are broadly similar: the shortening side of the simulated force-velocity relation follows the testing data, while the simulations of the lengthening side produce much less force than the test data. There are larger differences between the models when comparing the submaximal force-velocity relation of each model to Joyce and Rack [61]: the MAT_156, as expected, has a maximum shortening velocity of 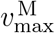(A); the EHTMM has a maximum shortening velocity that is slow compared to the experimental data (B); while the VEXAT model comes close to matching the experimental data (C). It should be noted that neither the EHTMM nor the VEXAT models were fitted to submaximal shortening data. In the time-domain all three models have similar RMSE values while shortening (D,E,F) and lengthening (G,H, and I), however once the length change ceases the VEXAT model’s force profile follows the experimental data [51] more closely than the other models (with the exception of D.). None of the models develops the amounts force-depression or force-enhancement contained in the experimental data set.

**Figure 5:**
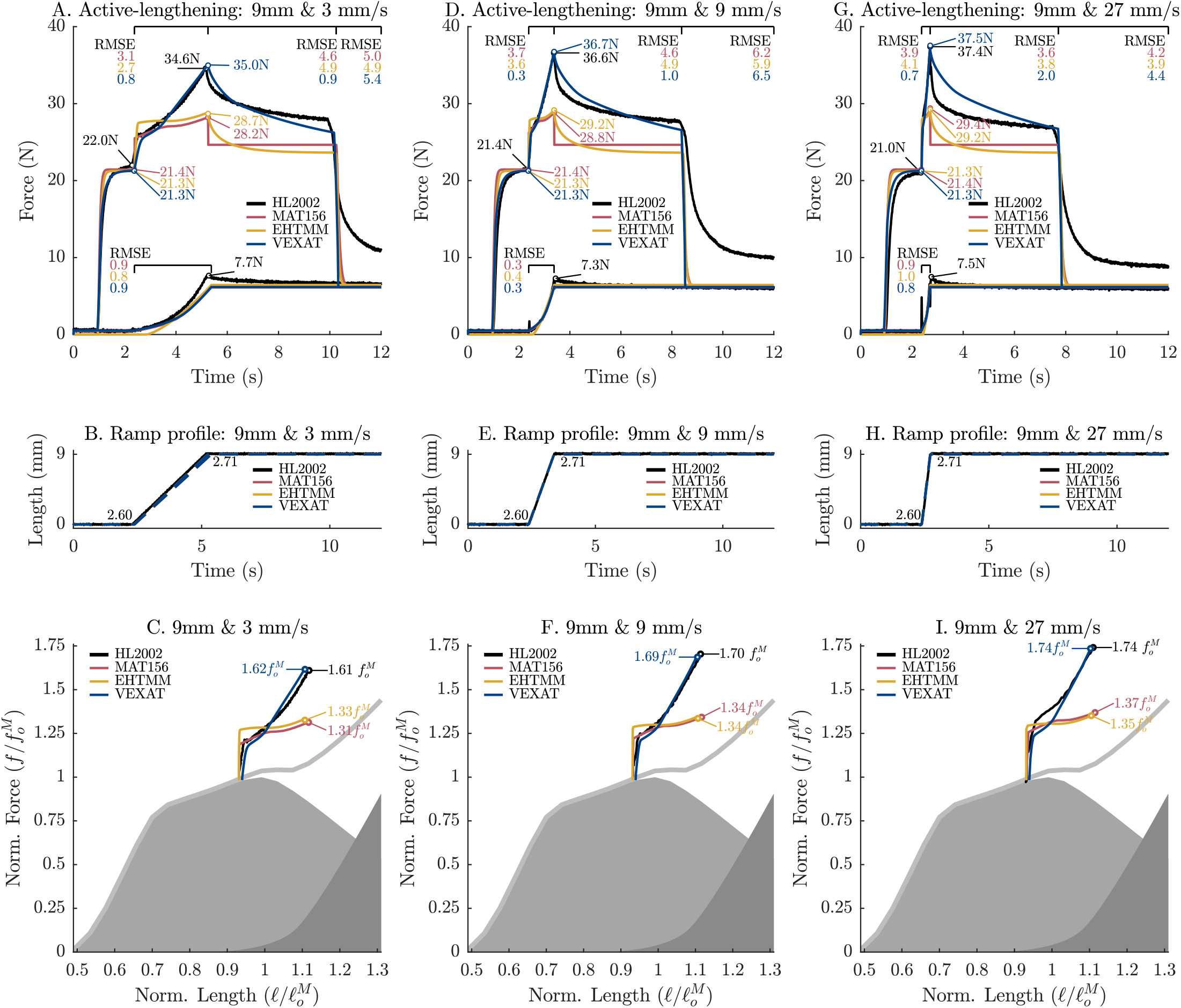
When the active lengthening experiments of Herzog and Leonard [11] are simulated using the HL02 model variant, the VEXAT model is able to reproduce the peak in force during the ramp (A, D, and G) and approximate the decrease in force following the ramp. The MAT_156 and EHTMM models consistently underestimate the experimental data. None of the models is able to produce the passive force enhancement present in the experimental data (A, D, and G): after deactivation, the tension of each model returns to passive values while the tension of the cat soleus remains elevated. Note that the VEXAT model’s titin element was fitted to the 9mm s^−1^ (D and E) trial, and so the 3mm s^−1^ (A and B) and 27mm s^−1^ (G and H) are test data.

**Figure 6:**
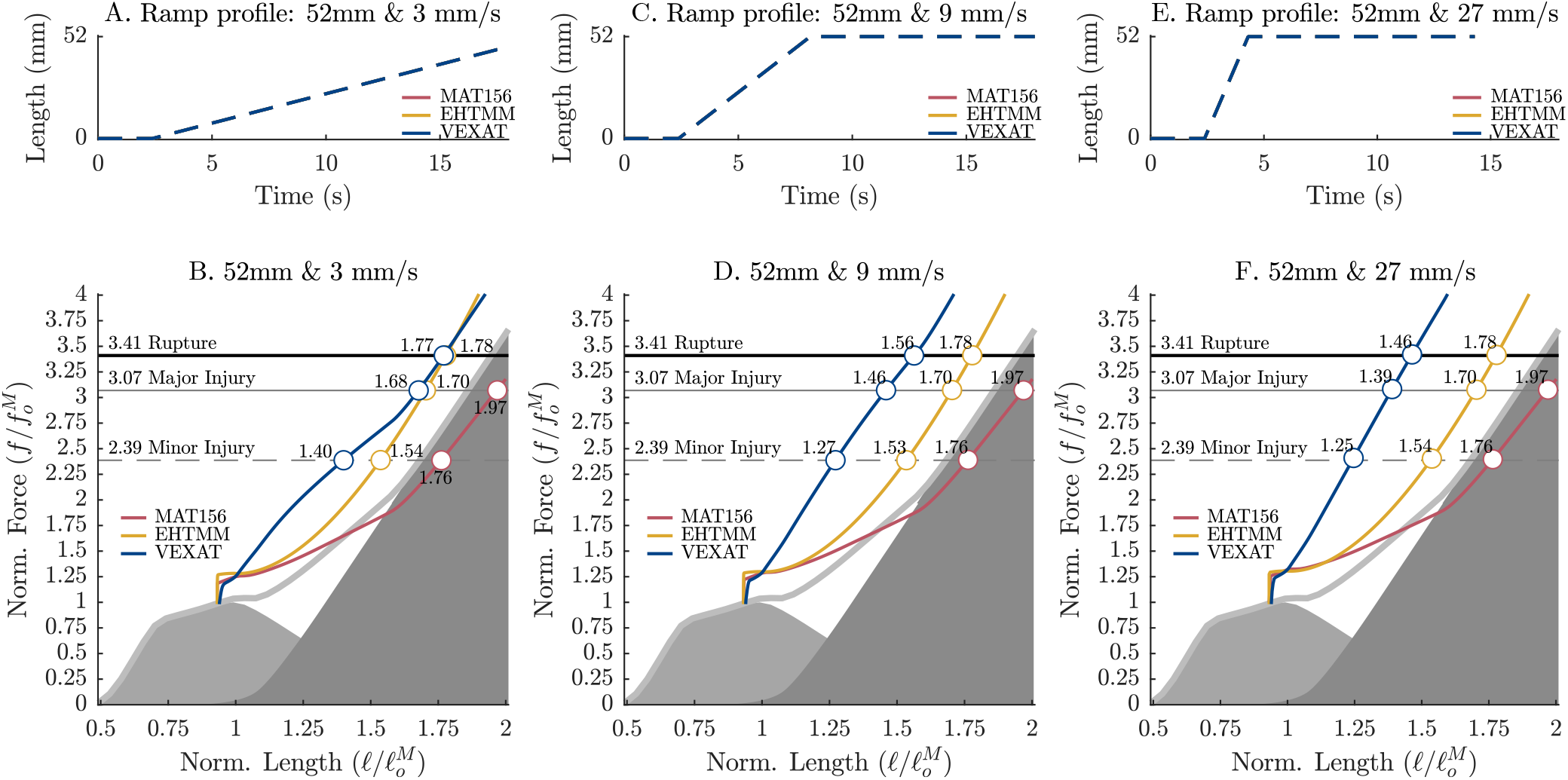
When the active-lengthening simulations (Fig. 5) are extended from 9mm to 52mm each of the models develops tension sufficient to pass through the mild, major, and rupture thresholds of active-lengthening injury [30], [62] though each model passes through these thresholds at different lengths. The VEXAT model passes through these thresholds at the shortest lengths of all three models because the active titin element allows it to develop active force even as 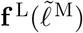 goes to zero [33], mimicking a surprising property of muscle [12]. Since the titin-actin bond of the VEXAT model is an activation dependent damper, velocity matters: during the 3mm s^−1^ trial (A) the VEXAT model reaches each injury threshold at longer lengths than the 9mm s^−1^ (B) and 27mm s^−1^ (C) trials. The MAT_156 and EHTMM models, in contrast, pass through the thresholds of injury at nearly the same lengths regardless of velocity because these models can only generate passive force beyond actin-myosin overlap. Note that the MAT_156 develops less passive force than the reference areas in grey (the VEXAT’s active and passive force-length curves) because the MAT_156 has a rigid tendon, and so, its CE has been made to have the same compliance of the CE and tendon of the other models.

**Figure 7:**
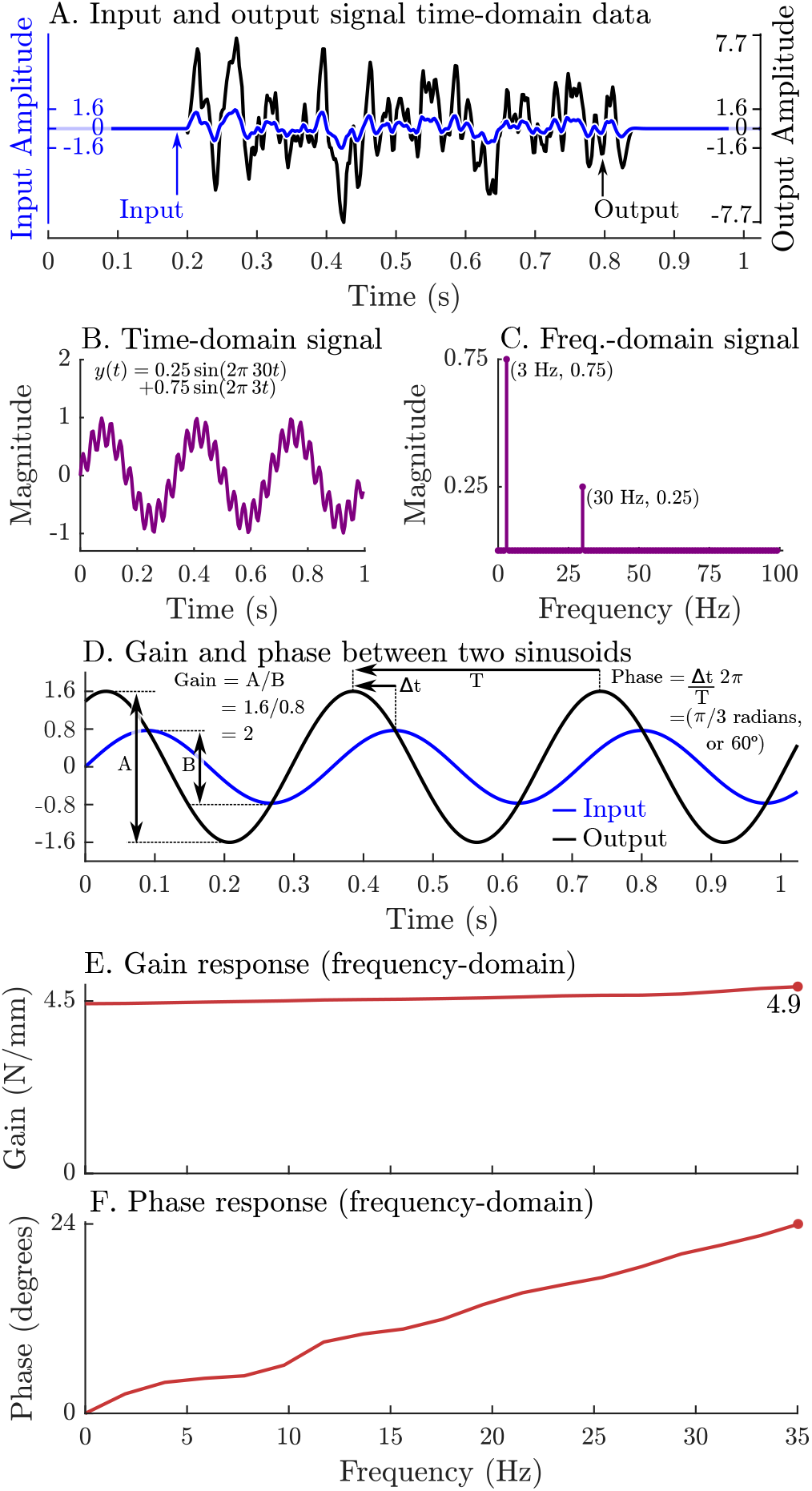
System identification methods can be used to identify a network of spring-dampers that best fits the response of muscle provided it can be treated as a linear time-invariant system. This process begins by constructing a stochastic bandwidth-limited length-change signal in the time-domain (A, blue line). Next, these length changes are applied to a muscle that is held at a constant nominal length and under constant stimulation (A, black line). These signals can be transformed from the time-domain (B) into an equivalent series of scaled and shifted sinusoids in the frequency-domain (C). The frequency-domain representation of both the input length-change and output force-response of muscle can be used to measure the relative amplitude (gain) and timing (phase) of the two signals (D). The pattern of gain (E) and phase (F) across a band-width of frequencies is often sufficient to identify (in this case) a network of spring-dampers that best fit the data.

**Figure 8:**
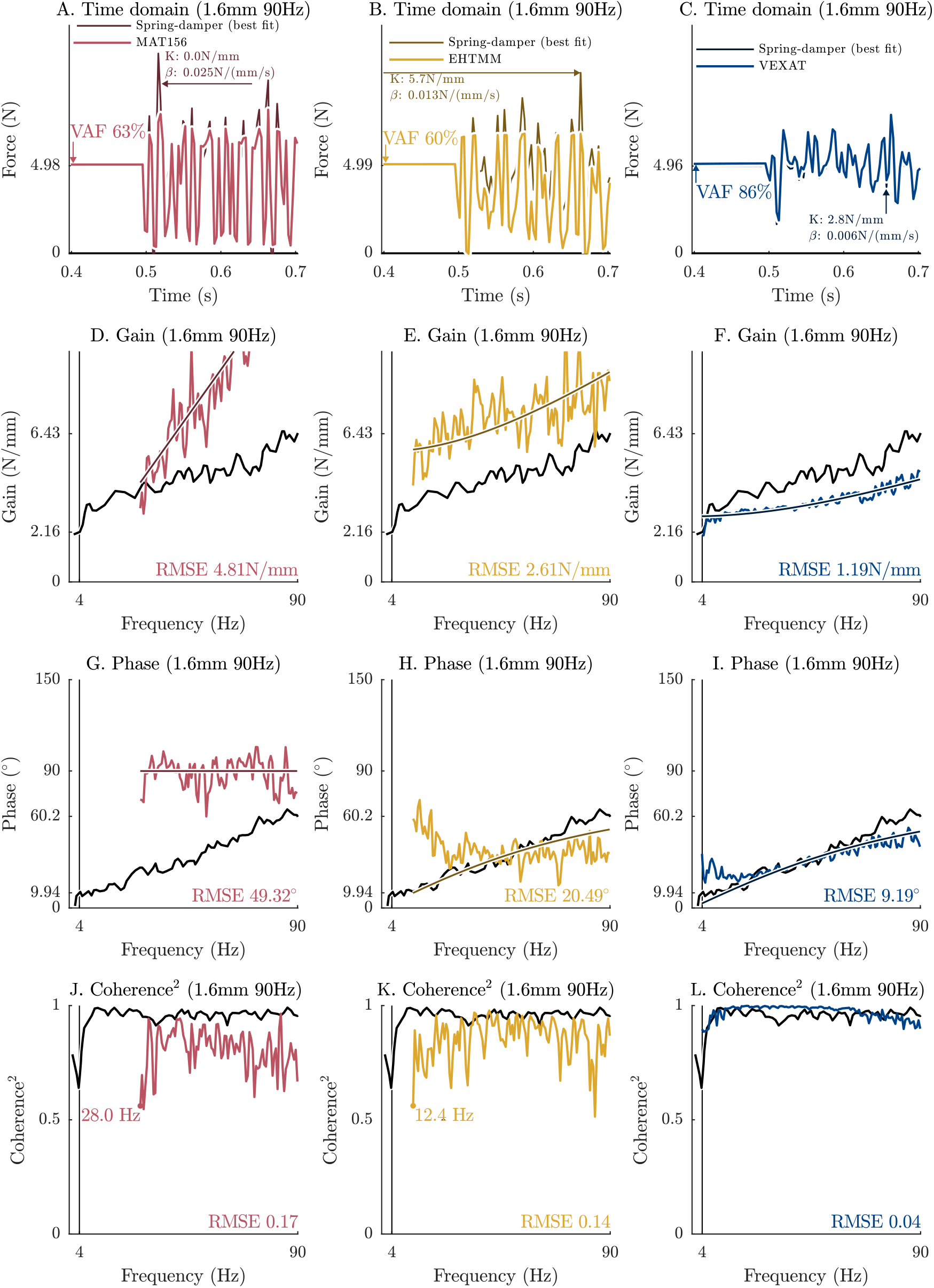
When the K3 variant is used to simulate Kirsch et al.’s [14] 1.6mm − 90Hz experiment there are marked differences between each model. In the time-domain the VAF (A, B, and C) shows that the MAT_156 and EHTMM have VAF values that are below the range of 78-99% reported by Kirsch et al. [14], while the VEXAT (C) model has a VAF that is within this range. Similarly, in the frequency domain both the gain-response (D, E, and F) and phase-responses (G, H, and I) of the MAT_156 and EHTMM deviate more from the experimental data [14] than the VEXAT model. We have taken care only analyze data for which the coherence-squared (a measure of linearity) exceeds 0.67 (J, K, and L) to be consistent with Kirsch et al.’s implied threshold ([14], Figure 3). While both the VEXAT model (L) and Kirsch et al.’s data [14] maintain a coherence-squared above the threshold for all frequencies above 4Hz, neither the MAT_156 (J) nor EHTMM (K) can meet this threshold at such low frequencies.

The maximally activated force-velocity trials show that all three have similar force-velocity relations (Fig. 4 A,B, and C), produce comparable forces during the length change (Fig. 4 D-I), though with some differences after the length-change has ended. The concentric side of the force-velocity relation of each model is similar to the measurements of Scott et al. [48], Brown et al. [49], and Joyce and Rack [61] while the eccentric side of the force-velocity relation is weaker than the datasets. In the time-domain, all three models show similar RMSE values during shortening (Fig. 4D-F), and lengthening (Fig. 4G-I). After active lengthening (Fig. 4G-I) the VEXAT model has a lower RMSE than either the MAT_156 or EHTMM due to the prolonged force-enhancement caused by the titin element. None of the models have the prolonged force-depression (Fig. 4D-F), nor the sustained force-enhancement (Fig. 4G-I) reported in Herzog and Leonard’s 1997 [51] measurements.

While concentric side of the force-velocity relation are similar between all experimental data sets, there are marked differences between the eccentric side of the force-velocity relation between the testing data sets [48], [49], [61] and the simulated models of which have been fitted (Fig. 1E) to the data of Herzog and Leonard 1997 [51]. Scott et al. [48] (Figure 6) provides a reason that might explain this difference: when force-velocity measurements are made at longer lengths force-enhancement increases. This may explain the difference in force enhancement between the data sets since Herzog and Leonard’s experiments [51] were performed at an ankle angle of 80^°^ (pg 866 paragraph 1 of [51]) which corresponds to a length estimated by our model to 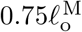 (when fully activated) while the measurements of Scott et al. [48] were made at 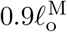 (pg 218 paragraph 1), Brown et al. [49] measured at 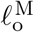 (pg 224 paragraph 1), and Joyce and Rack [61] report making measurements at an ankle angle of 70^°^ which Scott et al. [48] reports is equivalent to 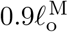.

To evaluate the sub-maximal force-velocity relations of each model, we repeat this entire set of simulations but with the excitation of each model set so that an isometric tension of 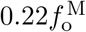 is developed prior to shortening to match one of the submaximal trials from Joyce and Rack [61]. Next, we extract the force-velocity relation from these submaximal simulations, and compare it to the sub-maximal force-velocity relation measured by Joyce and Rack [61] from an in-situ cat soleus. We have specifically chosen to simulate the sub-maximal trial that begins with a tension of 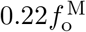 because the measurements of Joyce and Rack [61] show that at this tension the maximum contraction velocity is reduced from 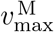 to 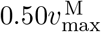, where we have identified 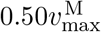 by fitting Hill’s force-velocity hyperbola [44] to the data.

The simulated submaximal shortening trials show 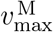 of the MAT_156 model (Fig. 4A) is unaffected by the reduced activation while both the EHTMM (Fig. 4B) and VEXAT (Fig. 4C) models have reduced contraction velocities. The submaximal contraction velocity of the VEXAT model 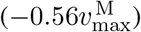 is slightly faster than Joyce and Rack’s data [61] 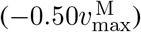 while the EHTMM 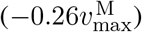 slower. None of the models follows the eccentric-branch of the submaximal force-velocity relation: as with the maximal contraction trials, the simulated submaximal trials level off during lengthening, while the measurements of Joyce and Rack [61] show that the force enhancement continues to increase with the rate of lengthening.

### 3.4 Active lengthening on the descending limb

Higher forces are generated when muscle is actively lengthened on the descending limb [11] than on the ascending limb [51]. This phenomena has long been of interest to muscle physiologists because on the descending limb the value of 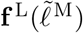 is *decreasing* during active lengthening, and yet the muscle is able to develop *higher* active forces. Muscle models also have had difficulty simulating active-lengthening on the descending limb since the active force of most models is proportional to 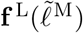, and so, decreases as the muscle is lengthened beyond 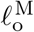. Since higher forces are generated when active muscle is lengthened on the descending limb, this phenomena is also of concern for simulations of injury: as tension continues to increase the muscle will be at increasing risk of injury [30], while at the same time its enhanced forces may prevent injury to other tissues by limiting the range-of-motion of a joint.

We examine the forces developed by the MAT_156, EHTMM, and VEXAT models during active lengthening by simulating experiments of Herzog and Leonard 2002 [11] in which an in-situ cat soleus is actively lengthened on the descending limb by 3, 6, and 9mm. To evaluate the accuracy of each model, we compare its peak force during lengthening to the experimental data [11] as well as compute the RMSE during three phases of the experiment: during the length change, after the length change, and finally after the muscle has been deactivated. For these simulations we make use of the HL02 parameters (Table 2) which have been fit to the passive force-length, active force-length, and force-velocity data that is embedded in the time-series data of Herzog and Leonard (see Figures 7A-C of [11]). Since Herzog and Leonard’s 2002 experiment [11] is well below the threshold of injury [30], [62], we also simulate the forces that are developed when the muscles are stretched by 52mm and compare the forces developed to the thresholds of active-lengthening injury [30], [62]. Unfortunately, we cannot directly replicate Hasselman et al.’s experiments [30] because the data needed to fit the models to Hasselman et al.’s specimens are not reported. As a result, we cannot compare the forces developed during injury to experimental data but can only make a comparison between the models.

The VEXAT model more accurately reproduces the force-profiles of the in-situ cat soleus during and after the 3mm s^−1^ (Fig. 5A), 9mm s^−1^ (Fig. 5D), and 27mm s^−1^(Fig. 5G) than either the MAT_156 or EHTMM models (see Appendix C Figs. 10 and 11 for the 6mm and 3mm trials). Once the model is deactivated, however, all of the models produce comparable forces and fail to produce the passive force-enhancement that is present in the experimental data [11]. When the ramp force profile is resolved in a normalized force-length space, the two-phase nature of the force enhancement is visible across each of the trials (Fig. 5C, F, and I): initially force develops rapidly during the ramp up to a force of 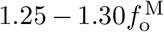 is reached, afterwards force continues to increase but at a lower rate. While all three models show the initial rapid force development, only the VEXAT model’s tension follows the experimental data [11] and continues to increase during the active-lengthening trial (Fig. 5C, F, and I).

**Figure 9:**
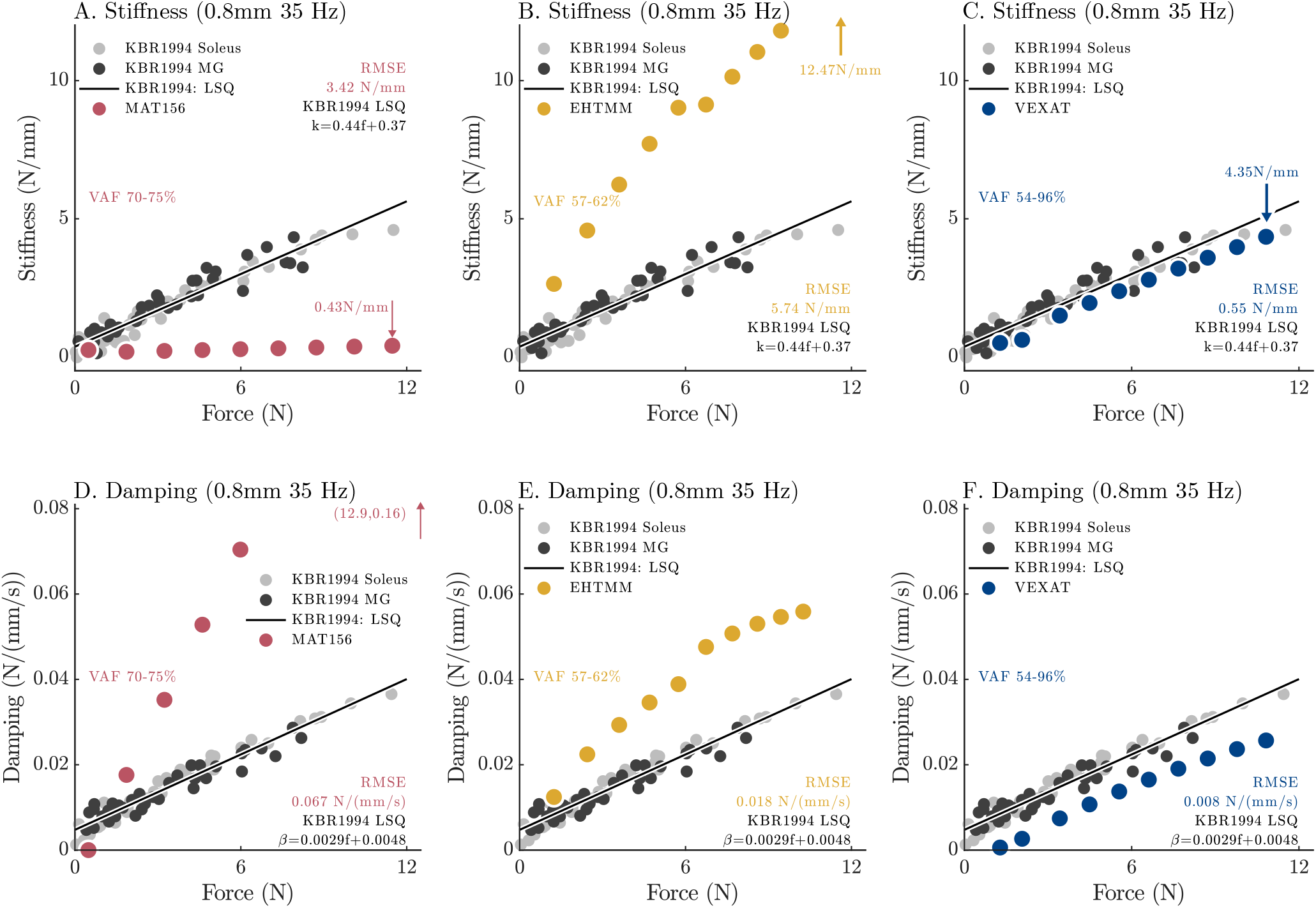
When Kirsch et al. [14] repeatedly applied perturbations (0.8mm − 35Hz) across a range of nominal forces (but with the same nominal length 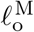) they observe that the stiffness (A, B, and C) and damping (D, E, and F) coefficients of best fit vary linearly with active force (see Figure 12 of [14]). Simulating this experiment using the K12 variant of each model shows that each model has distinct changes in stiffness and damping with nominal force. The MAT_156 model has a very low stiffness does not change with active force (A), while its damping increases more rapidly than the data [14] as active force increases (D). In contrast, both the stiffness (B) and damping (E) of the EHTMM are much larger than the data. The VEXAT model closely follows the stiffness (C) and damping (F) data.

**Figure 10:**
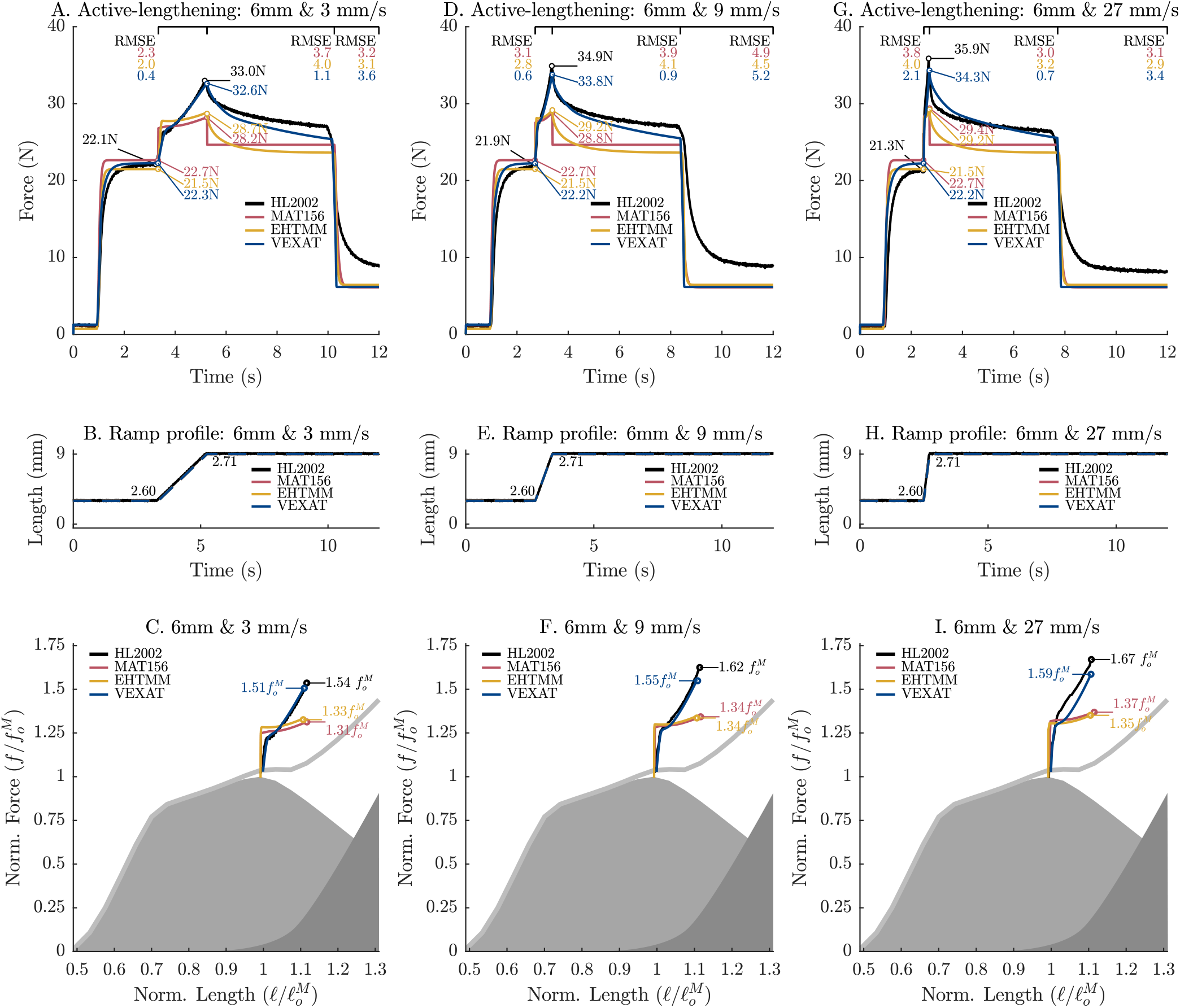
Herzog and Leonard [11] studied the effect of length change independently of the final length by starting the ramp 3mm longer but finishing at the same 9mm from the reference length for a total length change of 6mm (B,E, and H). The 6mm stretch produces lower peak forces than the 9mm stretch, a pattern that is replicated by the VEXAT model in both the time-series data (A, D, and G) and in the force-length space (C, F, and I). In contrast, both the MAT_156 and EHTMM produce the same peak forces (compare A, D and G to Fig. 4A, D and G) during the 6mm stretch as during the 9mm stretch. While none of the models develop passive force enhancement, the cat soleus [11] develops less passive force enhancement during the 6mm stretch than the 9mm. Note that the VEXAT’s titin model remain fitted to the 9mm − 9mm s^−1^ trial (Fig. 4D), and so, every trial pictured here can be considered testing data.

**Figure 11:**
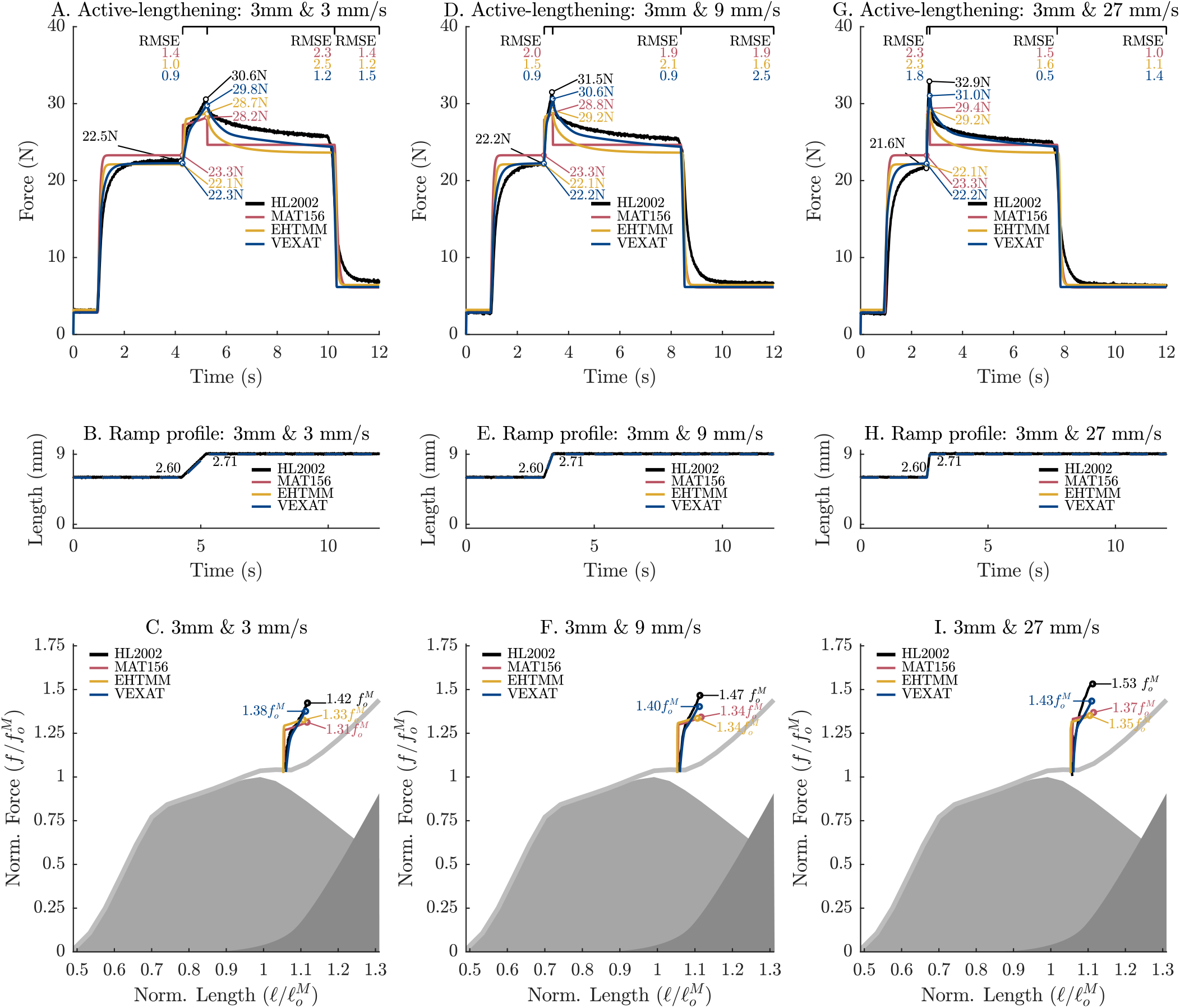
When the length change is reduced from 9mm (Fig. 4), to 6mm (Fig. 10), and finally to 3mm it is clear that the peak tension of both the cat soleus [11] and the VEXAT model vary together, producing lower peak forces as the length change is reduced. As before, both the MAT_156 and EHTMM produce the same peak forces independent of the size of the length change. As a result, the peak forces in both the time-domain (A, D, and G) and force-length space (C, F, and I) are quite similar during the 3mm length change. In addition, the cat soleus [11] produces virtually no passive force enhancement during the 3mm trial, and so, the models and the experimental data produce similar force at the end of the trial (see A, D, and G at second 12.0).

**Figure 12:**
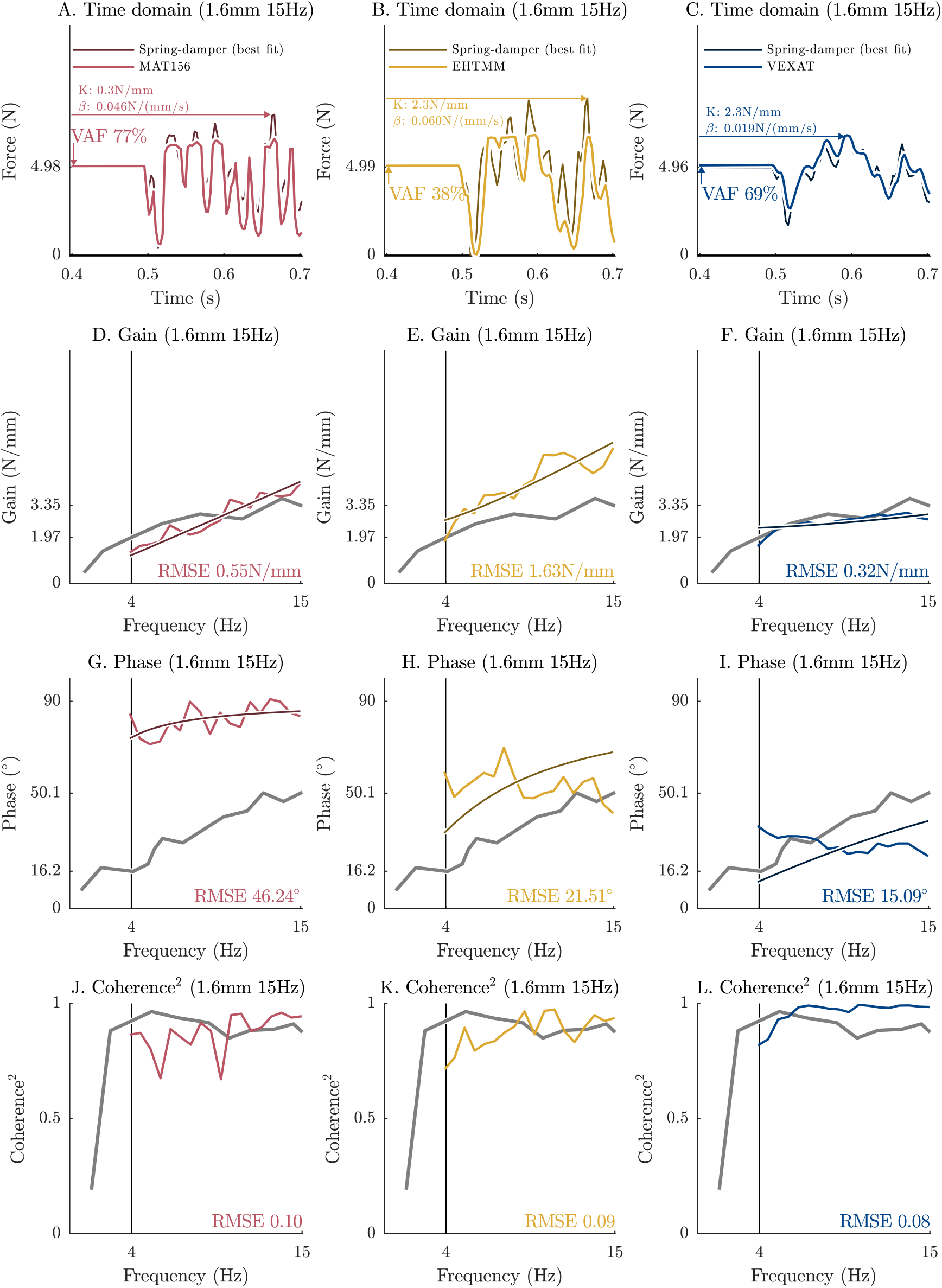
The response of the models to the 1.6mm − 15Hz perturbation differs with the response to the 1.6mm − 90Hz. In the time domain, the MAT_156’s VAF has improved (A), the EHTMM’s VAF has declined a lot (B), and the VEXAT’s VAF has declined a little. In the frequency-domain, the largest differences in comparison to the 1.6mm − 15Hz perturbation are: the accuracy of the gain response of both the MAT_156 and VEXAT have improved, as have the coherence-squared values of the MAT_156 and EHTMM models.

The VEXAT model is able to develop enhanced forces during active lengthening due to the active-titin element (Fig. 2E and F). When activated, a point within the VEXAT model’s titin segment becomes viscously bound to actin (Fig. 2E and F). As a result, the length of the proximal titin segment is approximately constant while the distal titin segment bears most of the strain and produces enhanced forces (Fig. 1F). After the ramp completes, the enhanced tension developed in titin’s distal segment relaxes as the viscous titin-actin bond slowly slides in response to the force imbalance until the enhanced force has completely dissipated (Fig. 6A, D and G). The passive force enhancement that is present in the experimental data [11] suggests that (using the VEXAT model to interpret the data) the force imbalance between the proximal and distal segments of titin should not completely dissipate. Both the MAT_156 and EHTMM models also develop enhanced forces during active lengthening due to the force-velocity and passive-force length relations though these mechanisms alone are insufficient to produce the enhanced forces present in the experimental data (Fig. 5A, D, and G). The force contribution of titin is even more prominent in fiber-level experiments of stretch-shortening [63] and extreme active lengthening [12] to the point of fiber-rupture.

Titin’s simulated force contribution becomes more pronounced when the active-length change is increased to cause injury. When the length change is extended from 9mm to 52mm at 3, 9, and 27mm s^−1^ (Fig. 6A, B, and C) the VEXAT model passes through the thresholds of injury before the EHTMM and MAT_156 models (Fig. 6D, E, and F) at each speed. During the 3mm s^−1^ trial the titin-actin bond has enough time to slip, and so, the VEXAT model passes through the thresholds for major injury and rupture at nearly the same normalized lengths as the EHTMM (Fig. 6A). The 9 and 27mm s^−1^ are quick enough that the titin-actin bond stays nearly fixed in place and, as a result, the VEXAT model passes through all injury thresholds at shorter normalized lengths than either the EHTMM or MAT_156 models (Fig. 6B, and C). Since the the CE of the MAT_156 has the lowest stiffness^11^, and lacks a titin element, it passes through the thresholds for injury at much longer normalized lengths than either model or the reference force-length curves (Fig. 6, reference curves in grey). The EHTMM passes through the thresholds of injury at shorter normalized lengths than the reference force-length curves (Fig. 6) because its passive force-length relation follows a power function whereas the VEXAT and MAT_156 models have a passive force-length relation that eventually becomes linear.

The difference in force development between the models during long active stretches can affect musculoskeletal simulations of injury. The larger forces developed by the VEXAT model can have two consequences: first, the VEXAT model will pass through the thresholds of injury at lower strains and become injured more quickly; second, the enhanced forces developed by the VEXAT model may protect the tissues of the joint it crosses by reducing the movement of the joint. The amount of force enhancement provided by the VEXAT model will vary with the titin isoform of the muscle and the stiffness of the ECM: shorter isoforms of titin will produce larger forces than longer isoforms (making the active titin force-length relation stiffer in Fig. 1F), while the difference between active and passive force-development will decrease as the ECM becomes stiffer (making both the active and passive titin force-length relations less stiff in Fig. 1F). In these simulations, we have used titin parameters from a human soleus titin [64] which has a long titin isoform, and the average of the titin and ECM contributions (56% ECM and 44% titin [33]) measured by Prado et al. [65] from rabbit skeletal muscle. Shorter isoforms of titin would be stiffer than the long isoform of titin we modelled [33]. Since Prado et al.’s measurements [65] of the relative contribution of titin and the ECM to the passive force-length relation are unique, we cannot know at this point in time if the relative contributions of titin and the ECM that we are using is appropriate for human skeletal muscle.

### 3.5 Active impedance of muscle

The active impedance of muscle increases linearly with active tension [14], a property that is exploited by the central-nervous-system (CNS) when learning new movements [66], to interact with mechanically unstable environments [67], and to reduce noise [68]. Muscular impedance is likely also important to accurately simulate the response of the body to vibration and ultimately to estimate vibration discomfort and motion sickness [5]. Since active muscular impedance can be represented as a stiff spring in parallel with a light damper [14] muscle impedance also contributes to the increase of force that is observed [11] during active lengthening.

Active muscle impedance [14] differs from short-range stiffness [69]. Rack and Westbury [69] coined the term short-range stiffness to describe a specific observation: during sufficiently small and rapid changes in length the change in force measured in active muscle is linear and independent of velocity. The stiffness, in short-range stiffness, is the ratio of force-change to length-change [69] during these small rapid length changes. In contrast, the impedance of muscle applies to the case when the changes in length and muscle force can be accurately reproduced using a linear time-invariant (LTI) system. LTI systems in the mechanical domain can include springs and dampers which produce force responses that are velocity dependent, and so, apply to a larger range of perturbations than short-range stiffness. The work of Kirsch et al. [14] showed that muscle under constant activation responds like a spring-damper in parallel to perturbations across a variety of bandwidths^12^ and amplitudes^13^. At frequencies lower than 4Hz, Kirsch et al. (see Figure 3B, [14]) found that the linear association between the length change and force output decreased — as quantified by the coherence-squared between the input and output ([70], pg. 137) — indicating that it was no longer appropriate to approximate the response of the muscle as an LTI system.

Approximating muscle as an LTI system makes its possible to identify an underlying set of equations and parameters that best fit the response of muscle over a bandwidth of frequencies. Kirsch et al.’s [14] experiments began by applying a small amplitude stochastic signal to vibrate the length of the active muscle causing it to generate a corresponding force response. Next Kirsch et al. [14] applied system identification methods to identify an LTI system that best captured how the muscle transformed the length changes into force changes during the experiments. To create the stochastic input signals Kirsch et al. [14] created a pseudorandom sequences of numbers between *±*1, filtered the signal using a second-order Butterworth filter (with −3dB frequencies of 15Hz, 35Hz, and 90Hz), and scaled the result (*±*0.4mm, *±*0.8mm, and *±*1.6mm) to the desired amplitude (Fig. 7A^14^ in blue). Next, the input and output signals are transformed into equivalent signals in the frequency-domain using a Fourier transform [71]. A Fourier transform [71] decomposes time-domain signals (Fig. 7B) into an equivalent series of sinusoids (Fig. 7C) that vary in frequency, scale, and phase but when summed together produce the original time-domain signal. As long as the muscle behaves like an LTI system there is a linear relationship between the input and output signals in the frequency-domain: the output will consist of the same set of sinusoids as the input except each sinusoid may have had its amplitude (gain) and phase-altered (Fig. 7D). The gain and phase-response (collectively known as the frequency-response) of an LTI system describes how the system transforms an input sinusoid to an output sinusoid (Fig. 7E and 7F show that a 1mm 35Hz input sinusoid will be transformed by the active muscle into a 4.9N, 35Hz output sinusoid that with a phase shift of 25^°^ relative to the input). Kirsch et al. [14] used the pattern of phase and gain responses across a broad range of frequencies to identify that a parallel spring-damper fits the response of muscle under constant activation. As the methods required to do this analysis are involved, we refer the curious reader to additional reference material ([33], Section 3.1 and Appendix D) and source code^15^ for further information.

To evaluate the impedance of the models we simulate the experiments of Kirsch et al. [14] and compare the time-domain response, frequency-domain response, and the impedance-force relation to the data of Kirsch et al. [14]. In the time-domain, Kirsch et al. [14] note that a fitted spring-damper will have a variance-accounted-for (VAF)

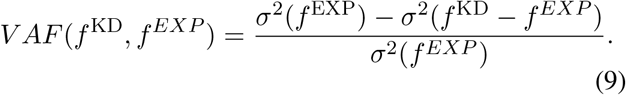

of between 78 − 99% for the cat soleus (60 trials) and medial gastroc (50 trials). We evaluate the time-domain response of each of the models by fitting a spring-damper to the response of each model and evaluate the VAF in the time domain to the *±*1.6mm−15Hz and to the *±*1.6mm−90Hz length change trials ([14], Figure 3). Using the response to the same *±*1.6mm − 15Hz and *±*1.6mm − 90Hz trials we evaluate the frequency-domain response by computing the RMSE between each model’s response and Kirsch et al.’s data ([14], Figure 3) of the phase-response and gain-response. For these simulations we use the K12 model variant (Sec. 3.1) in which the viscoelasticity of the VEXAT’s XE has been fitted to Kirsch et al.’s [14] Figure 3. When evaluating the frequency-domain response, we consider only the data above 4Hz and with a coherence-squared value of above 0.67 to be consistent with Kirsch et al. [14] (see the coherence-squared plot in Figure 3 of [14]). Next, we measure the response of each model to a 15Hz − 0.8mm perturbation as the active force developed by the model is linearly increased from 1 − 12N across a series of 10 trials. Model variant K3 (Sec. 3.1) is used for these simulations, where Figure 12 from Kirsch et al. [14] has been used to fit the viscoelasticity of the VEXAT’s XE. We fit a parallel spring-damper to each model’s frequency-response and compare how stiffness and damping vary with active force in comparison to Kirsch et al.’s data [14].

In the time-domain and frequency-domain the accuracy of each model differs depending on whether the *±*1.6mm − 90Hz or the *±*1.6mm − 15Hz perturbation is applied. In response to the *±*1.6mm − 90Hz perturbation, the VAF of the VEXAT model (86%, Fig. 8C) outperforms both the MAT_156 (63%, Fig. 8A) and EHTMM (60%, Fig. 8B) models. In the frequency domain, the RMSE of the VEXAT model is lower in both gain and phase (1.19N mm^−1^ and 9.19^°^, Fig. 8F) than either the MAT_156 (4.81N mm^−1^ and 49.32^°^, Fig. 8D) or EHTMM models (2.61N mm^−1^ and 20.49^°^, Fig. 8E). While VEXAT model’s coherence-squared values (Fig. 8L) remained well above the threshold of 0.67 at frequencies of 4Hz and higher (Fig. 8L), the lowest frequencies analyzed had to be raised for both the MAT_156 (28Hz, Fig. 8J) and EHTMM (12.4Hz, Fig. 8K) models meet the coherence-squared threshold.

Although the accuracy of each model’s response to the *±*1.6mm − 15Hz trial (Appendix D, Fig. 12) differ, when ranked by accuracy the result is similar to the *±*1.6mm − 90Hz trial. In the frequency-domain, the VEXAT model has a lower RMSE (0.32N mm^−1^ and 15^°^) with Kirsch et al.’s data [14] (Figure 3 [14]) than either the MAT_156 (0.55N mm^−1^ and 46.24^°^) or EHTMM (1.63N mm^−1^ and 22^°^) models. However, the response of the MAT_156 to the *±*1.6mm − 15Hz trial has a higher VAF (77%) than either the EHTMM (38%) or VEXAT (69%) models. All three models have sufficiently high coherence-squared values so that all data between 4 − 15Hz is included in the analysis.

The impedance-force relation of the VEXAT model (Fig. 9C and F) is similar to Kirsch et al.’s data [14] while the impedance-force relations of the MAT_156 (Fig. 9A and D) and EHTMM (Fig. 9B and E) differ. Since the length of the muscle is 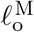 on average (where 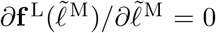), the stiffness of the MAT_156 is close to zero as would be expected from the derivative of Eqn. 3. The damping of the MAT_156, in contrast, increases with force at four times rate of Kirsch et al.’s data [14]. In contrast, the EHTMM’s response is quite different from the MAT_156. The elastic tendon and modified formulation of the EHTMM allow it to produce stiffness (Fig. 9B) and damping (Fig. 9E) responses that are larger than Kirsch et al.’s data [14]. These differences show up clearly in the RMSE stiffness and damping values from the VEXAT (0.55N mm^−1^ and 0.008N*/*(mm*/*s)), EHTMM (5.74N mm^−1^ and 0.018N*/*(mm*/*s)), and MAT_156 (3.42N mm^−1^ and 0.067N*/*(mm*/*s)) models (Fig. 9).

While the results we have found here differ strongly between the models, there is reason to expect that these results are sensitive to both the nominal length of the CE and the perturbation. The MAT_156 is a rigid tendon Hill-type muscle model, and as such, the active stiffness of this model depends on the nominal length of the CE: on the ascending limb a rigid-tendon Hill-type muscle model will have positive stiffness, at the optimal CE length the stiffness will go to zero, while on the descending limb the stiffness can become negative. The addition of an elastic tendon is likely the factor that gives the EHTMM an improved response in comparison to the MAT_156, as this pattern has also been observed in between other rigid-tendon and elastic-tendon Hill-type muscle models [33] (see Figure 7). The stiffness and damping coefficients of the VEXAT model will also be affected by the nominal length, since the 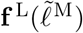 relation is multiplicative with the force developed by the XE in Eqn. 6.

Kirsch et al. [14] also observed that the stiffness and damping of best fit varies with both the frequency and amplitude of the perturbation (see Figure 3, 9, and 10 of [14]). While it is not yet clear what mechanism is responsible for this shift, there is a chance that this phenomena is tied to the cycling rate of cross-bridges: the 90Hz length perturbation is likely close to the cross-bridge cycling rate^16^, while 15Hz length perturbation is probably slower. At this point in time it is not clear what mechanism is responsible for this sensitivity to the perturbation, and so, it’s unlikely that any of the models evaluated would display the same pattern.

## 4 Discussion

Simulating injury using digital HBM’s is complex because of the wide variety of factors that can affect the calculated risk of injury. During a vehicle collision [75], athletic injury [6], [7], or in response to vibration [5], the body’s musculature may have time to activate, alter the movements of the body, and change the risk of injury. In this work, we have evaluated the accuracy of three different muscle models in LS-DYNA by simulating laboratory experiments that examine the force-length-velocity relations during maximal and submaximal activation, the response of muscle to active-lengthening, and the frequency-response of muscle. We have chosen to use the FE code LS-DYNA for our benchmark because LS-DYNA is frequently used to simulate injury sustained as a result of vehicle collisions [1], [2] and sporting accidents [7].

Our benchmark simulations are necessarily limited by the experimental data available on passive and active-lengthening injury from the muscle physiology literature. Passive and active-lengthening injuries have been measured in rabbit muscles [30], [31] and used by Nö lle et al. [62] to define the thresholds of passive and active-lengthening injury which we use in this work (Fig. 6). Unfortunately the works of Noonan et al. [62] and Hasselman et al. [30] do not contain the information needed to accurately fit a model and simulate the experiments, and so, we are left without an experimental reference for the simulations of active-lengthening injury that appear in Sec. 3.4. Even if the works of Noonan et al. [62] and Hasselman et al. [30] could be simulated, these studies may not be a good reference for the lengthening-injury characteristics of human skeletal muscle since Persad et al. ([76], Figure 6) recently illustrated that whole muscle in rabbits is far stiffer than whole muscle in humans. Due to the limited data on length-injury our benchmark can only make relative comparisons between models.

Although the experiments that measure the frequency-response of muscle [14] are more amenable to simulation than the lengthening injury experiments [30], [62], there are still a number of important experimental gaps that remain to be filled. Kirsch et al. [14] measured the frequency-response of cat soleus and medial gastrocnemius at the optimal fiber length, while the frequency-response of the ascending and descending limbs of the force-length relation have yet to be measured. Sugi and Tsuchiya [77] did measure the gain of frog skeletal muscle at a specific frequency during both shortening and lengthening, but did not measure the corresponding phase-shift. While the measurements of Kirsch et al. [14] are invaluable, there are still many open questions in regards to the frequency-response of muscle, and a sparse amount of experimental data in the literature.

The results of our benchmark simulations complement and extend prior work of Kleinbach et al. [32]. Kleinbach et al. [32] evaluated the activation dynamics, the force-length relation, and the concentric-force-velocity (quick-release) relation of the EHTMM and MAT_156^17^ models using data from a piglet plantarflexors [27] (force-length-velocity), cat soleus [78] (activation dynamics), and rat gastrocnemius [79] (activation dynamics). Briefly, Kleinbach et al. [32] showed that the EHTMM closely followed the experimental force-length data, more accurately captured the data from the quick-release experiment [27] than the MAT_156, and found Hatze’s [80] activation dynamics models to be more accurate than Zajac’s [45]. We have found that the force-length relation of the EHTMM closely matched the fitting data set [11], [51] (Fig. 1C) but deviated from the testing data set [21], [48], [50] at short CE lengths, and during submaximal activation (Fig. 3E) similar to the other models. In contrast to Kleinbach et al. [32], our simulations of the isokinetic force-velocity experiments using a cat soleus [51] found that the EHTMM and MAT_156 produce similar results (Fig. 4), though the EHTMM does have a lower RMSE than the MAT_156 during the ramp trial. The difference between the EHTMM and the MAT_156 may have been more pronounced during Kleinbach et al.’s [32] simulations because the tendon-to-CE length ratio is higher for the piglet plantarflexor 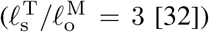 than for a cat soleus 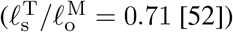. We have not included activation dynamics in the benchmark, as was done by Kleinbach et al. [32], but instead have simulated experiments in which activation is held constant so that our results do not depend on the activation model.

The benchmark simulations in this work also complement and extend our previous work [33]. Here we have evaluated the force-length (Fig. 3) and force-velocity (Fig. 4) relations using a broader set of experimental data than our previous work [33], and across both maximal and submaximal activation. While our previous work also includes simulations of active lengthening [33], here we have simulated a greater selection of the trials (Figs. 5, 10, 11) measured by in Herzog and Leonard [11], and simulated active lengthening injury (Fig. 6). Finally, across all benchmark simulations we have evaluated the MAT_156 [35], EHTMM [27], [32], [36], and the Fortran implementation of the VEXAT model, none of which were considered in our previous work [33]. What is similar between this benchmark and our previous work [33] is that we compared the VEXAT model against a Hill-type muscle model: previously we evaluated a damped-equilibrium Hill model [47] while here we have focused on the MAT_156 and EHTMM models. Even though the mathematical formulations of the damped-equilibrium [47], MAT_156 [35], and EHTMM [27], [32], [36] are substantially different, when simulated, these formulations share some of the same characteristics: tension is underestimated during active-lengthening on the descending limb; both the MAT_156 and rigid-tendon damped-equilibrium model [33] are too compliant and too damped at 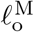 (compare Fig. 9A and D to Figure 7C and D from [47]); and while both the EHTMM and elastic-tendon damped equilibrium model [33] have positive stiffness and damping at 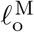 these values are large in comparison to Kirsch et al.’s [14] data (compare Fig. 9B and E to Figure 7B and D from [33]). Despite the differences in formulation, the active-lengthening and frequency-response of the rigid-tendon Hill models are similar, as are the responses of elastic-tendon Hill models.

In this work, we have evaluated three muscle models that can be used in LS-DYNA: the MAT_156 model, a Hill-type CE model; the EHTMM model, a Hill-type muscle model that includes a viscoelastic tendon; and the VEXAT model, a model that includes an active-titin element and a viscoelastic CE. While all three models performed similarly in the force-length and *ascending limb* force-velocity benchmark simulations, we found substantial differences during the *descending limb* active-lengthening, and frequency-response benchmark simulations. Consistent with previous work, Hill-type muscle models that lack an active-titin element will underestimate the force developed by the CE during active lengthening during modest (Fig. 5) and long stretches (Fig. 6). Muscles that underestimate active-lengthening forces may produce HBM’s that have a higher risk of injury than reality: compliant muscles will allow joints to bend excessively, perhaps injuring ligaments, that would otherwise be protected by stiffer muscles. Similarly, Hill-type muscle models have a frequency-response that differs substantially from experimental data [14]. When the response of a muscle model to vibration differs widely from experimental data so too will an HBM that uses these muscle models. While the VEXAT model performs better than either the MAT_156 or EHTMM during simulations of active-lengthening or in response to vibration, our work indicates that the performance of the model during submaximal force-length (Fig. 3H) and force-velocity simulations can be improved (Fig. 4C).

## 5 Conclusions

While the MAT_156, EHTMM, and VEXAT muscle models in LS-DYNA have comparable force-length and force-velocity relations, these models differ during active-lengthening on the descending limb and in response to vibration. During active-lengthening on the descending limb the VEXAT model’s titin-element allows it to produce enhanced forces similar to biological muscle, while the force respone of both the MAT_156 and EHTMM is too weak. In response to vibration the VEXAT model has a force profile that closely resembles a spring-damper in parallel, similar to biological muscle, while the MAT_156 and EHTMM are too damped.

## Acknowledgements

Financial support is gratefully acknowledged from the Deutsche Forschungsgemeinschaft (DFG, German Research Foundation) under Germany’s Excellence Strategy (EXC 2075 – 390740016) through the Stuttgart Center for Simulation Science (SimTech). We would also like to acknowledge Lennart Nölle, Maria Hammer, and Isabell Wochner from the Institute for Modelling and Simulation of Biomechanical Systems (IMSB) at the University of Stuttgart for the assistance they provided with the EHTMM model and LS-DYNA.

## A Fitting the passive and active force-length relations

The passive and active force-length relations of the VEXAT and EHTMM models are fit to experimental cat soleus data from the ascending [51] and descending limb [11] of the force length relation using a custom made fitting routine. First, we digitized the length change and recorded forces of the passive (δ*𝓁* ^PE*^, *f* ^PE*^) and active (*δ𝓁* ^L*^, *f* ^L*^) isometric data points just prior to the ramp movement from the ascending [51], and descending limb [11] of the force-length relation. Note that the asterix in *f* ^PE*^ and *f* ^L*^ are being used to denote a parameters fitted to a specific study rather than using more cumbersome notation such as ^97^*f* ^PE^ and ^97^*f* ^L^ for [51], and ^02^*f* ^PE^ and ^02^*f* ^L^ for [11]. In both studies, there are some parameters that are uncertain or unreported: the optimal CE length 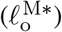, the maximum isometric force 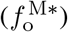, and the path length of the muscle that corresponds to the reference length of 0mm (*𝓁* ^R*^). Accordingly, our vector of optimization parameters x includes the 3 experimental parameters from each study 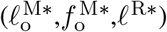 along with the parameters needed to shift (Δ^*^) and scale (*s*^*^) the passive-force-length relation of the VEXAT model to best fit the data.

No additional parameters are needed to fit the activeforce-length relation of the CE, nor the tendon force-length relation of the VEXAT model. The shape of the VEXAT model’s active-force-length curve has been made to follow the theoretical sarcomere-force-length relation proposed by Rassier et al. [59] which is depends on the length of the actin and myosin filaments (1.12*µ*m and 1.6*µ*m in cats). Preliminary simulations indicate that Rassier et al.’s [59] theoretical active force-length curve fits that of a cat soleus, though it should be noted that this is not true in general [58]. Only two parameters are needed to scale the normalized tendon force-length curve^18^ (Fig. 1B) to fit the data: the tendon slack length 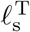, and the maximum stiffness of the tendon. Using a candidate value for 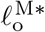 we solve for 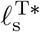 by assuming that the tendon-to-CE ratio as measured by Scott and Loeb [52] is maintained (27 mm of tendon to 38 mm of CE). Given a candidate value for 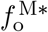 we can now scale the stiffness of the tendon force-length model such that it develops the same normalized tendon stiffness of 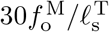 measured by Scott and Loeb [52].

The error of the **f** ^PE^ of the VEXAT model is evaluated first by using the bisection method to solve for the length of the CE that puts the passive CE and the tendon in a static force equilibrium

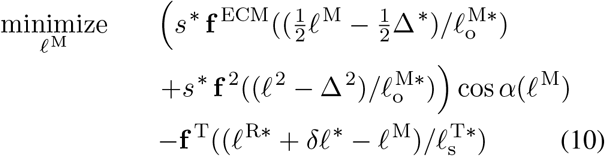

to mimic the experiment. At each iteration, we use Newton’s method to solve for the length of the proximal titin segment *𝓁* ^1^ that puts the proximal and distal titin segments in a passive force equilibrium

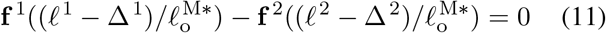

where

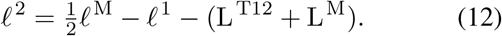

To ensure that the passive force-length relation of the model is adjusted by the desired amount Δ^*^ we also must shift the serially connected titin curves, which we do by distributing Δ^*^ across the proximal

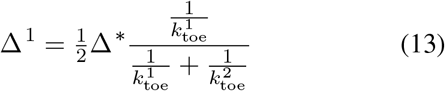

and distal

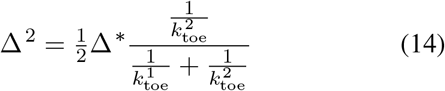

titin segments in proportion to the relative compliance of each segment. As with the tendon curve, the stiffness of both the proximal and distal titin curves varies nonlinearly up to a maximum stiffness of 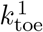 and 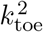.

Finally, the error of the model (*ϵ* ^PE^) is the difference in passive force developed by the model

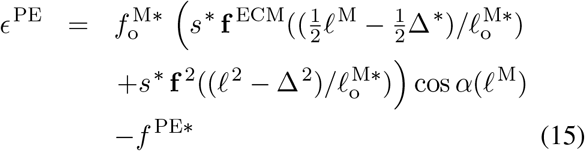

and the passive force (*f* ^PE*^) measured in the experiment. A similar procedure is used to evaluate the error of the active force developed by the model, where the bisection method is used to evaluate *𝓁* ^M^ that puts the active CE and the tendon in equilibrium

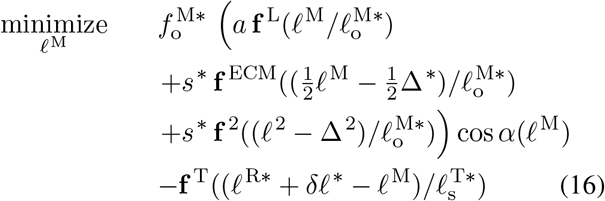

where *a* is the activation of the CE which is set to 1 for all of the active data. The active force error is the difference of the total isometric force produced by the model

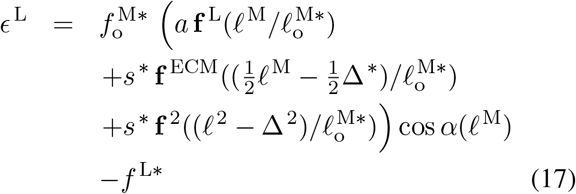

and the measured force (*f* ^L*^).

We use this approach to simultaneously solve for the parameters (see Tables 1A-C and 2A-C for the parameters of model variants HL97 and HL02 respectively) that minimize the active (Fig. 1C) and passive force-length (Fig. 1D) errors on the descending limb [11] using the Matlab [81] function *lsqnonlin*. The fitting procedure for the ascending limb data [51] is similar, though we restrict *s*^*^ to the value that best fits the descending limb data set [11] and only allow the optimization routine to shift **f** ^PE^ (Fig. 1D): there are too few passive data points in the ascending limb data set [51] to reliably fit both *s*^*^ and Δ^*^. The resulting fitted passive and active force length relations were numerically sampled and used to populate the tabular data that defines **f** ^PE^ and **f** ^L^ curves of the MAT_156.

The EHTMM is fit using a similar method though only the variables associated with the shape of **f** ^T^, **f** ^PE^, and **f** ^L^ were adjusted: the values of 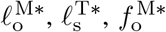 and 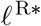 identified using the VEXAT model were used when fitting the EHTMM. First, the shape of the EHTMM’s **f** ^T^ was fit to the VEXAT’s **f** ^T^ (Fig. 1B) by varying a subset of parameters 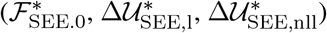 to minimize the sum of squared errors of the strain at 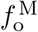, the stiffness at 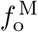, and the force developed in the middle 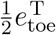 of the toe-region. Using the fitted tendon model, we simultaneously fit the variables that control the shape of the passive 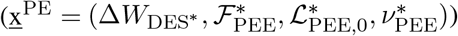 and active force-length 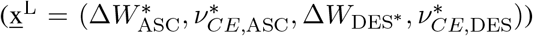 relations of the EHTMM to the passive and active data from the descending limb of the force-length relation [11]. As before, first we solve for *𝓁* ^M^ such that

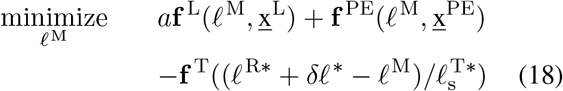

the tension developed by the CE and the tendon are equal under isometric conditions. When evaluating the error of the passive force length relation *a* = 0 in Eqn. 18 the error is evaluated as

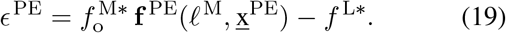

In addition, we also included two additional error terms from the fitted VEXAT **f** ^PE^: the force developed at 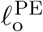 (where 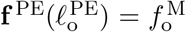), and the stiffness at 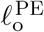. These extra points were added to ensure that the two models are similar when developing large passive forces. The error for the active isometric forces is evaluated as

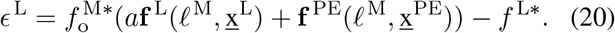

using the value of *𝓁* ^M^ that satisfies the force equilibrium in Eqn. 18 with *a* = 1. Using these error functions we solved for the parameters (see Tables 1A,F-H and 2A,F-H for the parameters of model variants HL97 and HL02 respectively) that simultaneously minimized the sum of squared errors across Eqns. 19 and 20 for the dataset on the descending limb [11] (Fig. 1C-D). As before, when solving the passive parameters x^PE^ for the ascending limb data set [51] we limited the optimization routine to shifting the passive curve that best fits the descending limb data [11] (Fig. 1D).

## B Fitting the force-velocity relation

With most of the architectural 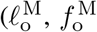 and 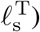, experimental (*𝓁* ^R*^), and force-length relations (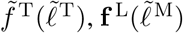 and 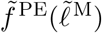) fitted we can now fit the force-velocity relation. To start, we digitize the following key points from Figure 1A of Herzog and Leonard 1997 [51]: the isometric force *f* ^M*^ developed at the final length of *𝓁* ^M*^ = 0mm, the forces 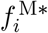 developed at the final ramp length of 0mm for all of the *i* = 1 … 5 shortening trials (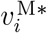 varies from −2.5 to −30 mm s^−1^), and the forces developed at the final ramp length of 0mm for all of the *i* = 6 … 10 lengthening (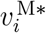 varies from 2.5 to 30 mm s^−1^) trials. Using these digitized points, we can transform this data into a series of discrete measurements that approximate the force-velocity relation using Eqn. 7 at the normalized velocities evaluated by Eqn. 8 which are in units of 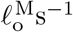. Although there is clearly some passive force being developed at *𝓁* ^M*^ (between 1-3N between *t* = 0 − 0.3s in Figure 1A of [51]) we ignore this passive component for two reasons: no measurement of this force is provided and it is small in comparison to *f* ^M*^ (37.5N).

Next, we fit the force-velocity relation of the VEXAT model so that its 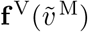 curve best fits the points 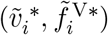. Since the experimental measurements of 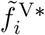 [51] are inline with the tendon, our first step is to estimate *𝓁* ^M^, *α*, and *v* ^M^ given 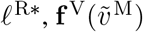. First, we are going to assume that the lengthening rate of the tendon is negligible (*v*^T^ ≈ 0), which is reasonable for muscle-tendon complexes in which 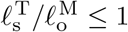 [47]. All model variants in this work have the same 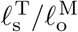 ratio of 0.71 (27mm*/*38mm = 0.71) as measured by Scott et al. [52]. Using this assumption, we can estimate the length of the tendon (ignoring damping) by inverting the force-length curve of the tendon

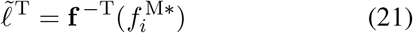

which allows us to solve for

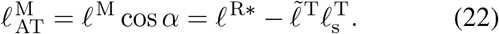

This assumption also allows us to relate

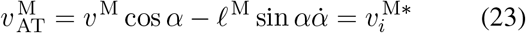

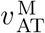 to 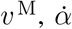 and 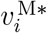. Since Eqn. 2 of the pennation model constrains the height of the CE to be constant we can solve for

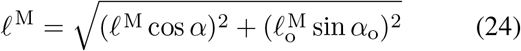

which allows us to solve for *α*

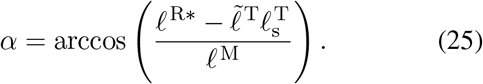

in Eqn. 22. By taking the derivative of Eqn. 2 we can solve for

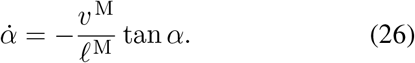

After substituting Eqn. 26 into Eqn. 23 we are left with

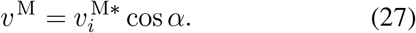

allowing us to evaluate

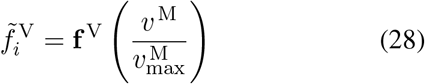

and calculate the error

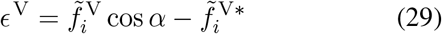

of the model’s force-velocity relation. By minimizing the sum of squared errors using the *lsqnonlin* function in Matlab [81] we arrive at values of 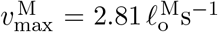 for HL97, and 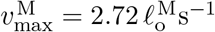 for HL02 variants of the VEXAT model (see Tables 1D and 2D for the parameters of model variants HL97 and HL02 respectively). The resulting **f** ^V^ of the VEXAT model fits the concentric data quite closely but deviates from the eccentric data points (Fig. 1E) with an overall root mean squared error (RMSE) of 0.0749. Although the eccentric side of the **f** ^V^ appears to be weak, the active-titin element of the VEXAT model will contribute additional tension that will be separately fitted at a later stage.

These values for 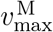 can be transformed to the non-pennated MAT_156 by noting that the CE length of MAT_156

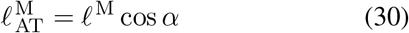

is the projection of the VEXAT model’s CE onto the direction. Taking a derivative we can solve for the rate of lengthening of the MAT_156 CE

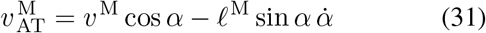

by substituting 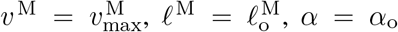 and evaluating 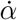 using Eqn. 26. This process results in values for 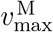 of for the MAT_156 model of 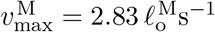 for HL97, and 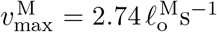 for HL02. The values for 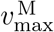 of both the VEXAT and MAT_156 models are very similar (see Tables 1E and 2E for the parameters of model variants HL97 and HL02 respectively) because *α*_o_ is small.

Defining the error function to fit the force-velocity relation of the EHTMM is less complicated than the VEXAT model because it is not pennated. Given a candidate set of parameters x = (B_rel,0_, F_e_, S_e_) we calculate the value of

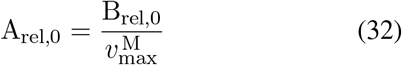

so that all three models share the same maximum shortening velocity of 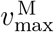. Now we can evaluate the error of the candidate **f** ^V^ as

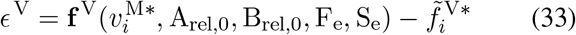

where **f** ^V^ is the force-velocity curve of the EHTMM where the concentric side is described in Günther et al. [27], and the eccentric side of the curve comes from Appendix A.1 of van Soest and Bobbert [82]. As before, we used Matlab’s [81] function *lsqnonlin* to minimize the sum of squared errors between the force-velocity relation of the EHTMM and the force-velocity data extracted from Figure 1A of Herzog and Leonard 1997 [51] for the HL97 and HL02 model variants (see Tables 1I and 2I for the parameters of model variants HL97 and HL02 respectively). The fitted **f** ^V^ of the EHTMM follows the data very closely for both the concentric and eccentric data points (Fig. 1E) as indicated by the low RMSE of 0.0255.

## C Additional active-lengthening simulations

## D Additional active impedance simulations

## Footnotes

1 The frequency-response refers to how the gain and phase of an input sinusoid are transformed by a system (muscle in this case) across a bandwidth of frequencies.

2 https://github.com/mjhmilla/Millard2024VEXATMuscleLSDYNA

3 https://github.com/mjhmilla/SingleMuscleSimulationsLSDYNA

4 All of the benchmarks make use of cat soleus which has a pennation angle of around 7^°^ [39].

5 The length of the CE at which 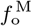 is developed during an isometric contraction.

6 The images of the MAT_156 and VEXAT models in Figure 2 have been used under the terms of the CC-BY license3 and have been modified from the original form [37]. The images in this figure are also licensed under the terms of the CC-BY licence3. A copy of the license can be found at https://creativecommons.org/licenses/by/4.0/legalcode

7 The impedance of a mechanical component is its stiffness and damping. The active impedance of muscle increases linearly with active force [14] and is referred to as the impedance-force relation in this work.

8 Which we normalize using 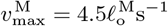 as reported on page 211 paragraph 3 of the results section [48]

9 Which we normalize using 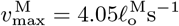 which is obtained using Equation 10 and the values of *b*_1_*/a*_1_ reported in Table 1 [49].

10 Which we normalize using 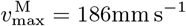 which we solved for by fitting Hill’s hyperbola [44] to the 35 impulses/second trial.

11 The MAT_156 has been fitted to have the same stiffness as the VEXAT model’s CE and tendon in series.

12 4 − 15 Hz, 4 − 35 Hz, and 4 − 90 Hz

13 *±*0.4mm, *±*0.8mm, *±*1.6mm and *±*6mm were evaluated which amounts to 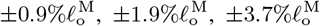 and 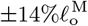 for a 42.9mm cat soleus

14 This figure is being used under the terms of the CC-BY license3 [33]. A copy of the license can be found at https://creativecommons.org/licenses/by/4.0/legalcode

15 See main SystemIdentificationExample.m in the eLife2023 branch of https://github.com/mjhmilla/Millard2023VexatMuscle

16 In-vitro measurements have been made of myosin filaments sliding at 3 − 4µm s^−1^ [72]. If each cross-bridge produces a step of 11nm [73] then we have a total of 273 − 364 cycles per second coming from the 98 [74] cross-bridges per half-myosin (assuming all cycles have the same step-lenth). Since the duty cycle ranges from 0.07 [73] under low-load up to 0.2-0.4 under isometric conditions we are left with a range of estimated cross-bridge cycling rates that vary between 19 − 146Hz.

17 The EHTMM was compared to the MAT_156 only during the quick-release experiments.

18 As is typical [45], the tendon force-length curve varies nonlinearly between strains of 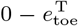 during which it develops forces between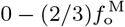. Strains greater than 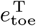 produce tendon forces that vary linearly with a stiffness of 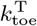.

